# A flexible high-throughput cultivation protocol to assess the response of individuals’ gut microbiota to diet-, drug-, and host-related factors

**DOI:** 10.1101/2023.07.11.548549

**Authors:** Janina N. Zünd, Serafina Plüss, Denisa Mujezinovic, Carmen Menzi, Philipp R. von Bieberstein, Tomas de Wouters, Christophe Lacroix, Gabriel E. Leventhal, Benoit Pugin

## Abstract

Anaerobic cultivation of fecal microbiota is a promising approach to investigate how gut microbial communities respond to specific intestinal conditions and perturbations. Here, we describe a practical protocol using 96-deepwell plates to cultivate stool-derived gut microbiota. Our protocol addresses key challenges in high-throughput culturing, including a thorough assessment of the impact of gas phase on medium chemistry, a modular medium preparation process to enable testing of several conditions in parallel, a medium formulation designed to maximize the compositional similarity of fecal cultures with the donor microbiota, and the creation of a step-by-step protocol detailing all practical procedures from material preparation to sample handling for analyses. Finally, we validated the protocol by demonstrating that cultivated fecal microbiota responded similarly to dietary fibers (resistant dextrin, soluble starch) and drugs (ciprofloxacin, 5-fluorouracil) as reported in vivo. This high-throughput cultivation protocol can facilitate culture-dependent studies, accelerate the discovery of gut microbiota-diet-drug-host interactions, and pave the way to personalized and microbiota-centered interventions.

**MOTIVATION:** The human gut microbiota is a complex ecosystem unique to each individual. The extent of this diversity has limited our capacity to fully comprehend microbial dynamics that apply to the entire human population. To probe the response of donor-specific microbial communities to intestinal conditions or perturbations, *in vitro* cultivation of stool-derived gut microbiota can be employed. However, cultivating gut microbiota under strictly anaerobic conditions is commonly performed using individual gas-tight tubes, which is a highly time-consuming strategy that limits the number of conditions and donor microbiota that can be tested in parallel. A flexible high-throughput protocol to cultivate and test donor-specific gut microbiota is therefore required. Hence, we developed a robust procedure for cultivating stool-derived microbiota (and pure gut microbial cultures) in 96-deepwell plates within an anaerobic chamber.

## INTRODUCTION

The human gut is inhabited by a diverse and dynamic assembly of microbes unique to each individual^1^. This microbial community contributes to key health functions such as nutrient metabolism, pathogen colonization resistance, maintenance of gut barrier integrity, and education of the host immune system, among others^2^. Despite advances in understanding the human gut microbiota’s diversity and functionality, the mechanisms governing ecological and physiological dynamics in response to specific intestinal conditions and perturbations remain poorly understood. Numerous studies have documented changes in the gut microbial community in response to dietary interventions^3–5^, antibiotic/non-antibiotic medications^6–8^, or alterations of the host physiology (*e.g.,* intestinal pH, oxidative stress, transit time, antimicrobial peptide production, and osmotic pressure)^9–11^. However, high inter- and intra-individual variability of the gut microbiota, combined with the complex interactions between diet-, drug-, and host-related factors *in vivo*, have limited our capacity to generalize microbial responses to a broader population^12^.

*In vitro* fecal cultivation holds promise for investigating the direct impact of relevant intestinal conditions on individuals’ gut microbiota. However, traditional cultivation of intestinal communities using anaerobic tube-based techniques (*e.g.,* Hungate tubes, Balch tubes or serum flasks)^13–15^ is time-consuming and has important limits on the number of donor microbiota and conditions that can be evaluated simultaneously. This is a major drawback as many responses and activities are specific to donor^16–19^, strain^20–23^, and physicochemical environment of the gut^9–11^. Cultivation with a higher throughput is thus essential to expand testing of conditions and our understanding of gut microbiota-diet-drug-host interactions and facilitate the development of personalized microbiota-centered interventions^4,24^. Although high-throughput cultivation in multi-well plates is not new^25–28^, several gaps remain unaddressed. The effect mediated by the anaerobic chamber gas phase on the chemistry of the growth medium and on microbial physiology has not been directly evaluated. A highly modular medium preparation process for simultaneously testing several culture conditions has yet to be developed. The formulation of a standardized growth medium mimicking the microbial composition of the donor feces in fecal cultures is still required. Lastly, a cultivation protocol detailing all technical procedures and steps with practical solutions is still lacking, which is crucial for ease of implementation and reproducibility in cultivation experiments.

In this study, we report the development and validation of a practical, flexible, and reproducible high-throughput protocol to cultivate stool-derived gut microbiota and pure intestinal microbes in 96-deepwell plates within an anaerobic chamber. This protocol builds upon a previous tube-based bacterial enrichment approach, where specific bacterial taxa were associated with the functional niches of the human gut^24^. Here, we transitioned the enrichment tube-based conditions to a format with higher throughput, and we expanded the applicability of the protocol for testing microbiota responses to a range of diets (*e.g.,* fibers, nitrogen sources, vitamins), drugs (*e.g.,* antibiotics and non-antibiotic medication), and host-related factors (*e.g.,* pH, redox potential, oxidative stress). We validated our protocol by showing similar *in vitro* responses to known dietary fibers (resistant dextrin, soluble starch) and drugs (ciprofloxacin, 5-fluorouracil) as reported *in vivo.* A comprehensive protocol detailing the necessary steps and procedures for material preparation, high-throughput cultivation, and sample handling for analyses is provided in **Supplementary file 1.**

## RESULTS

### Optimizing the procedures for high-throughput cultivation within the anaerobic chamber

We aimed to create a standardized high-throughput cultivation protocol that is compatible with standard anaerobic chamber setups and utilizes commonly available laboratory equipment. The first critical steps to transition our previous tube-based cultivation conditions^24^ to a high-throughput format included i) selecting a versatile and reusable multi-well vessel, ii) designing a robust yet modular medium preparation process, and iii) carefully assessing confounding factors associated with cultivation in the anaerobic chamber.

To replace the gas-tight rubber-stoppered tubes used with the gold-standard Hungate technique^14^, we opted for 96-deepwell plates combined with gas-permeable sealing membranes. Using 96-deepwell plates enables simultaneous testing of multiple conditions, with 60 wells allocated for experiments surrounded by 36 water-filled wells to minimize evaporation and edge effects^29^ (**Supplementary file 1, Step 4**). The working volume of 2 mL per well provides a sufficient amount of sample to conduct a range of analyses on the whole culture (*e.g.*, OD_600_, pH), the cell-free supernatant (*e.g.*, metabolites), and the cell pellet (*e.g*., DNA-based analyses) (**Supplementary file 1, Step 8**).

To prepare the medium and adjust the culture conditions, we designed a modular procedure in which an anaerobic basal medium solution (containing nitrogen (N)-sources, fatty acids, salts, buffers, trace elements, and heat-stable vitamins) can be flexibly complemented within the chamber with a carbon (C)-sources solution, or any other solution with specific components (heat stable/sensitive) to be tested (**Figure 1A; Supplementary file 1**). For the basal medium, we formulated bYCFA and mM2, which are modified versions (C-depleted, reduced N-sources) of the yeast extract casitone and fatty acids (YCFA) and the M2GSC media, respectively, two widely used media for cultivating gut microbes and microbiota^30,31^. Using such media with minimal nutrient availability allows identification of supplement-specific signals. In contrast to rumen fluid containing mM2 (30% v/v), bYCFA includes only readily-available compounds and thus facilitates accessibility and reduces batch-to-batch variability, as rumen fluid varies in composition and is difficult to procure.

**Figure 1:**
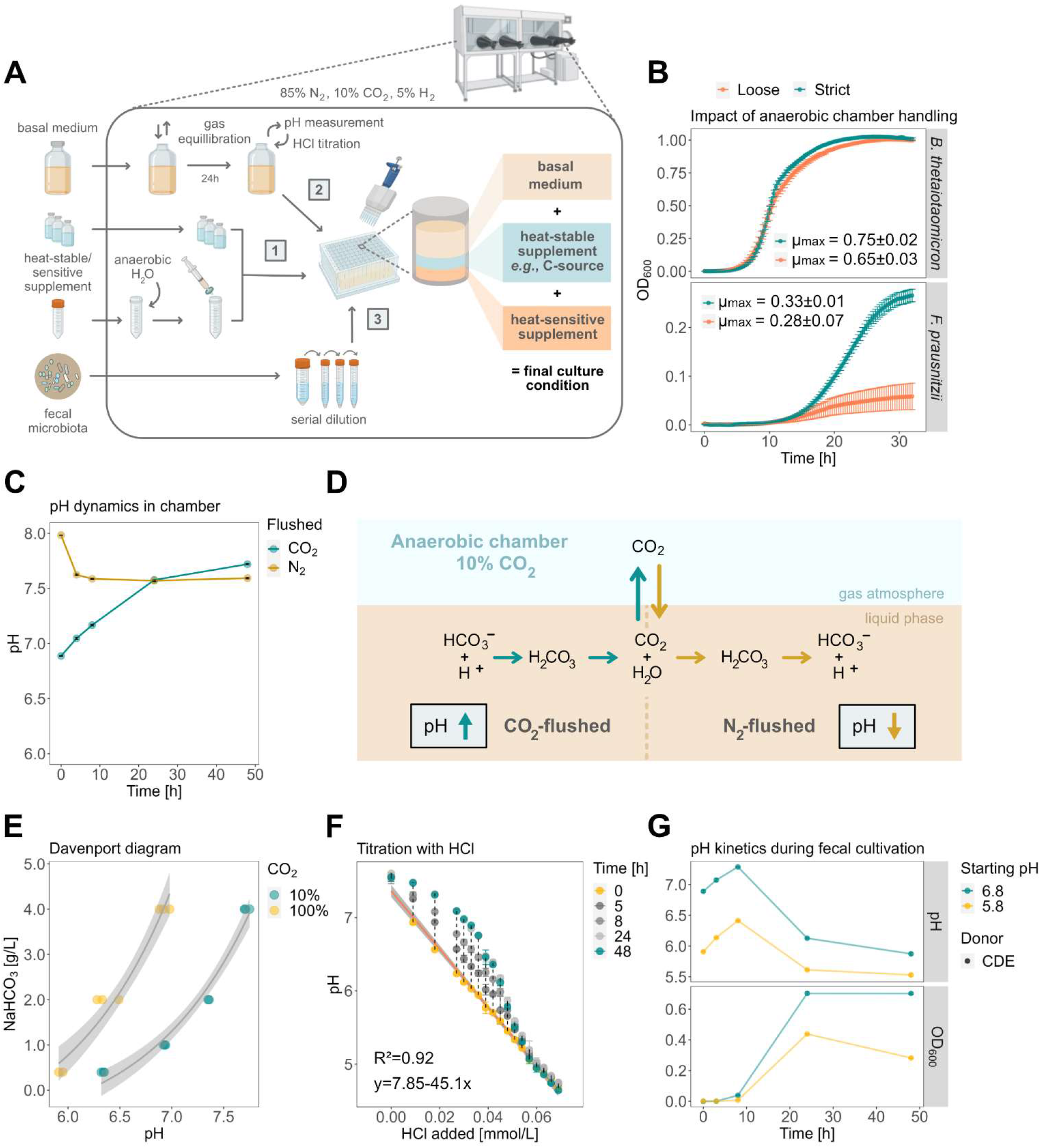
Overview of the high-throughput cultivation protocol and effect of the anaerobic chamber atmosphere on the medium’s chemistry and microbial physiology. A) Overview of the step-by-step cultivation protocol adapted for the anaerobic chamber setup. In a 96-deepwell plate, heat-stable/sensitive supplements (1) are mixed with HCl-titrated bYCFA (2) and inoculated with diluted feces (3). B) Growth kinetics (OD_600_) of *B. thetaiotaomicron* DSM 2079 and *F. prausnitzii* DSM 17677 in bYCFA (30 mM glucose, pH 6.5, 37°C, 32 h) in an anaerobic chamber that was handled “strictly” (turquoise dots; >2.5% H_2_) and “loosely” (coral dots; <1.5% H_2_). Each condition was tested in biological triplicates. C) Characterization of pH dynamics of CO_2_ and N_2_-flushed bYCFA (uninoculated) in a 96-deepwell plate in an anaerobic chamber. D) Impact of gas phase of the anaerobic chamber on the pH of basal medium previously flushed with CO_2_ or N_2_. E) Relationship between pH, aqueous NaHCO_3_ concentration, and gaseous CO_2_ levels (∼10% in the anaerobic chamber, ∼100% in gas-tight tube;) in bYCFA after 48 h equilibration. F) Change of pH over time upon titration of bYCFA with HCl, with subsequent pH increase over 48 h. The resulting titration curve (at time 0 h) can be used to adjust the pH to different values. G) pH and growth (OD_600_) during 48 h batch fecal cultivation in bYCFA (initial pH 6.8 and 5.8; containing 4 g/L NaHCO_3_ and 3 g/L resistant dextrin) titrated with HCl immediately before inoculation (donor CDE; 1% inoculation of 10^−4^ fecal dilution). Each condition was tested in technical triplicates.

An initial challenge when cultivating intestinal microbes within an anaerobic chamber is maintaining strict anaerobiosis. During plate-based cultivation, cultures are exposed to the chamber’s atmosphere, which can be contaminated with O_2_ that diffuses in or is introduced with consumables. Residual O_2_ is removed by reacting with H_2_ and the palladium catalyst, producing H_2_O. To show the impact of incomplete O_2_ removal on bacterial growth, we cultivated two intestinal bacteria, one less sensitive to O_2_ (*Bacteroides thetaiotaomicron* DSM 2079) and one highly sensitive to O_2_ (*Faecalibacterium prausnitzii* DSM 17677), in a “strict” and “loose” O_2_-handling regime. In the loose regime, the chamber was improperly prepared with non-regenerated catalysts and low H_2_ (<1.5%) in the atmosphere, whereas the strict regime had regenerated catalysts and the H_2_ level was >2.5%. The growth rate (μ_max_) of *B. thetaiotaomicron* was slightly but significantly higher in the strict compared to the loose regime (0.75±0.02 vs. 0.65±0.03 h^−1^; p<0.05). For *F. prausnitzii*, we found higher variation and lower μ_max_ when incubated in the loose regime (0.28±0.07 h^−1^) compared to the strict regime (0.33±0.01 h^−1^; **Figure 1B**; **Table S1**). This illustrates that strict preparation of the chamber (**Supplementary file 1, Step 2**) is crucial for reproducing anaerobic microbial growth.

A second challenge during cultivation in the anaerobic chamber is pH control, which is influenced by the equilibrium between gaseous and dissolved CO_2_, with the latter forming carbonic acid (H_2_CO_3_) in an aqueous solution^32^. When preparing the basal medium, 100% CO_2_ or N_2_ is flushed to remove traces of O_2_, resulting in dissolved CO_2_ levels that are out of equilibrium with the chamber atmosphere (10% CO_2_, 5% H_2_, and 85% N_2_). As a result, introducing CO_2_-flushed medium in the chamber led to a significant abiotic increase of pH (+0.16±0.05 after 4 h, and +0.83±0.03 after 48 h in bYCFA). In contrast, a rapid drop of pH was observed when introducing N_2_-flushed medium in the chamber (−0.36±0.02 after 4 h in bYCFA, **Figure 1C**; similar pH dynamics in mM2, **Figure S1**). These changes in pH reflect release of CO_2_ into the gas phase (CO_2_-flushed) or the solubilization of CO_2_ into the medium (N_2_-flushed) to equilibrate with the levels in the chamber’s atmosphere (**Figure 1D**). Regardless of the gas employed during media preparation, it is thus recommended to equilibrate the medium in the chamber for at least 24 h prior to the experiment (**Supplementary file 1, Step 3**). The pH at equilibrium depends on CO_2_ level in the atmosphere as well as the concentration of NaHCO_3_ in the medium, *i.e.,* the conjugate base of H_2_CO_3_ in the bicarbonate buffer system (Davenport diagram, **Figure 1E**). To establish a standard NaHCO_3_-containing basal medium that can be flexibly adjusted to a range of pH values, we chose a fixed concentration of 4 g/L NaHCO_3_ and opted to adjust pH within the chamber via HCl titration (**Supplementary file 1, Step 6**). This approach allows rapid adjustment of the initial pH for cultivation; however, there is a slow increase in pH over time (**Figure 1F**) as HCl titration shifts the bicarbonate buffer system towards CO_2_ that is gradually released again, resulting in alkalinization. The pH should thus be adjusted right before inoculation to maintain the initial pH in the desired range. To validate the feasibility of this approach, we titrated bYCFA with HCl (to pH 6.8 and pH 5.8) immediately before inoculation with fecal microbiota and incubated the cultures at 37°C for 48 h (**Figure 1G**). After an initial pH increase of +0.48±0.07 (pH 6.8) and +0.45±0.02 (pH 5.8) after 8 h, the pH decreased and reached final values of 5.90±0.04 (pH 6.8) and 5.51±0.04 (pH 5.8) after 48 h (**Figure 1G**). The degree of alkalinization resulting from the disrupted bicarbonate buffer was small compared to the acidification caused by microbial activity (*i.e.,* organic acid production), and the pH remained within the range of the bicarbonate buffering system (pH 5.1 to 7.1)^32^ throughout the cultivation. Therefore, HCl-titration was integrated into the protocol for establishing the final culture conditions. This step requires a pH meter in the chamber and using a pre-determined titration curve (**Supplementary file 1, Step 6; Figure 1F**) to rapidly set the initial pH to the desired values.

### Effect of basal medium and gas phase composition on the metabolism and community structure of fecal cultures

Having selected a high-throughput vessel and establishing a modular procedure for preparing the medium and adjusting culture conditions, we next aimed to evaluate whether the basal medium composition could influence fecal cultures’ metabolic activity and taxonomic composition. We compared bYCFA and mM2 media, with bYCFA considered a potential substitute for mM2 without rumen fluid. As an alternative to the growth factors in rumen fluid, bYCFA contains a mixture of hemin, vitamins and short-chain fatty acids (SCFA). Using both the plate-based and tube-based cultivation techniques, we cultured two fecal microbiota (48 h, 37°C) in bYCFA and mM2 supplemented with two different C-sources (resistant dextrin, soluble starch, or H_2_O as control). Our results showed that the concentration of total metabolites (R=1, **Figure S2A**) and individual metabolites (*i.e.,* SCFA and intermediate metabolites; R=0.97, **Figure S2B**) correlated for the two media. Similarly, the richness of fecal cultures determined via observed ASVs (alpha diversity) strongly correlated between the two basal media (R=0.94, **Figure S2C**). The community composition, assessed via Bray-Curtis distances (beta diversity), showed closely overlapping confidence ellipses when visualized with Principal Coordinate Analysis plot (PcoA plot; **Figure S2D**). Taken together, these data indicate that the effect mediated by the basal medium was minimal. Therefore, to ensure reproducibility and minimize batch-to-batch variability, we recommend using bYCFA as the standard basal medium for cultivating fecal microbiota (**Supplementary file 1**).

Next, we aimed to quantify to what degree the gas phase conditions of both cultivation techniques could impact fecal cultures, *i.e.,* an open system with gas-permeable 96-deepwell plates exposed to 10% CO_2_, 5% H_2_ and 85% N_2_ and a closed system with gas-tight tubes containing 100% CO_2_. In bYCFA (**Figure 2A**; mM2 in **Figure S3**), the concentrations of total metabolites (R=0.99, **Figure 2B**) and individual metabolites (R=0.97, **Figure 2C**) produced strongly correlated between the two cultivation techniques. The only noticeable difference was a lower accumulation of the intermediate metabolite formate in plates as compared to tubes when starch was added (**Figure 2C**). The richness of fecal cultures (observed ASVs) was similar or higher in plates than in tubes (**Figure 2D**). Permutational multivariate analysis (Adonis2) of Bray-Curtis distances revealed that cultivation technique had the smallest effect size (F) on community composition (F=3.9, p≤0.01), as highlighted by the closely overlapping confidence ellipses for the two cultivation techniques (**Figure 2E**). Variations were mostly explained by the donor microbiota (F=70.1, p<0.001), followed by the supplemented fiber (F=24.1, p≤0.001). Finally, specific taxa enriched with each fiber also correlated between plates and tubes (R=0.85, **Figure 2F**).

**Figure 2:**
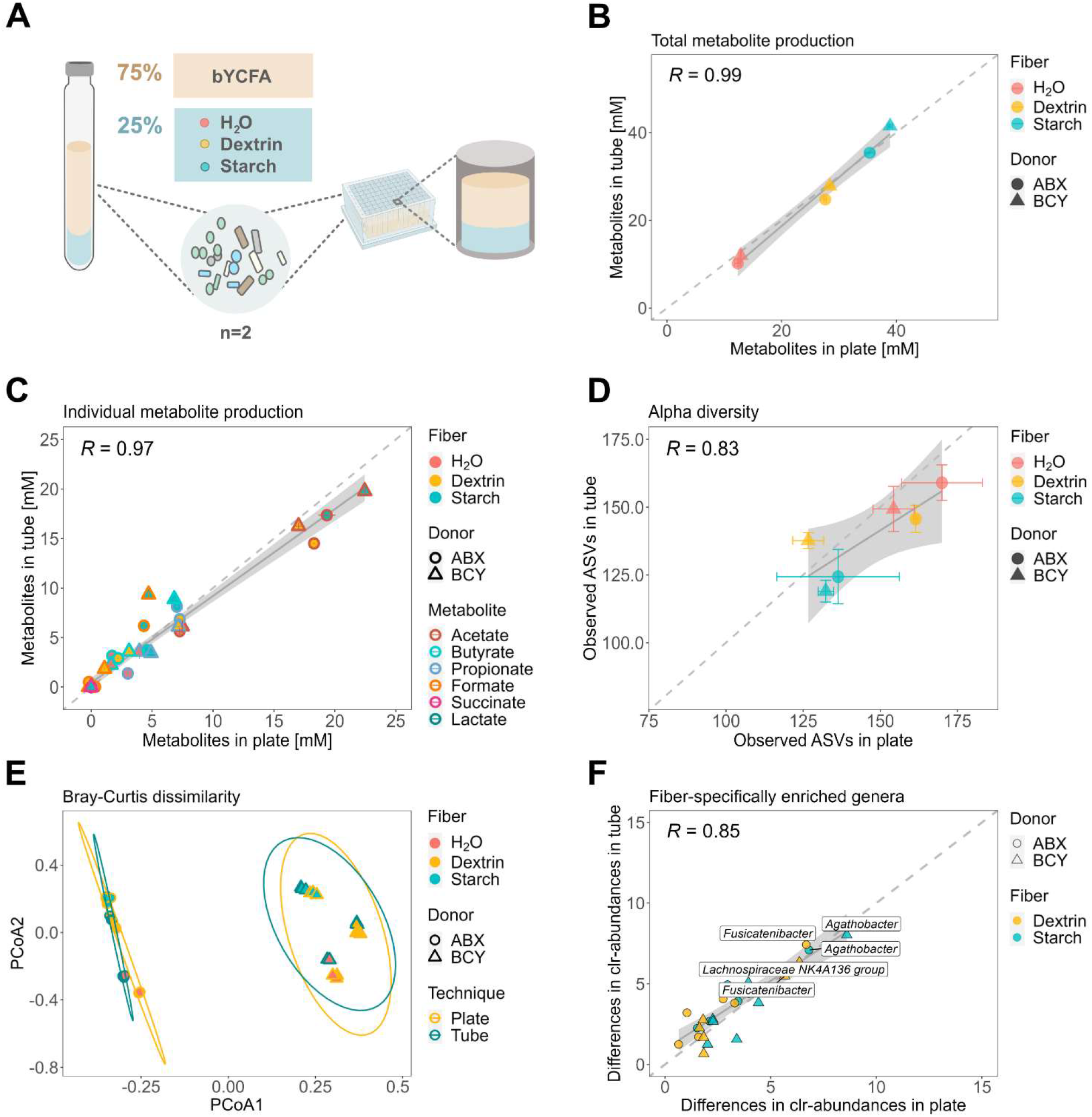
Impact of the cultivation technique (96-deepwell plates vs. gas-tight tubes) on the metabolism and composition of fecal cultures (donor ABX and BCY) cultivated in bYCFA for 48 h, 37°C, in the presence of resistant dextrin (3 g/L), soluble starch (3 g/L) or H_2_O as control (technical triplicates). **A)** Experimental setup for comparing the plate- and tube-based techniques. All analyses were performed after 48 h cultivation. **B)** Correlation of the total metabolite production. Total metabolites were calculated by summing the medium-corrected concentrations of organic acids (SCFA and intermediates). **C)** Correlation of individual metabolites. Points represent means of blank corrected concentrations and bars represent standard deviations. **D)** Correlation of observed ASVs. Points represent means and bars represent standard deviations. **E)** PCoA plot of Bray-Curtis distances with 95% confidence ellipses (calculated separately for each donor) for the two cultivation techniques. **F)** Median clr-differences of taxa (>1% abundance) that are significantly different between each fiber (dextrin, starch) and H_2_O (p ≤0.05). Labels indicate highly enriched taxa (clr-differences >5).

Though the fecal culture profiles were similar when grown using the two cultivation protocols, subtle differences in the metabolites produced were observed (*i.e.,* reduction of formate concentration both with bYCFA (**Figure 2C**) and mM2 (**Figure S3C**)). To identify whether these differences might be attributed to slight variations in the metabolic outputs of individual strains exposed to the chamber’s atmosphere, we cultured a panel of pure gut bacteria in bYCFA using both cultivation techniques (**Table S2**, **Figure S4A**). The selected bacteria covered a range of nutritional preferences (*e.g.,* primary degraders of complex fibers, utilizers of intermediate metabolites) and metabolic outputs (*e.g.,* acetate-, propionate- and butyrate-producers). All bacteria were able to grow in the anaerobic chamber using the plate-based technique, though six out of eight strains reached lower OD_600_ values after 48 h compared to the tubes (**Figure S4B**). Cultivation technique appeared to influence the carbohydrate metabolism of the tested strains, with all cultures exhibiting higher levels of end-metabolites (the SCFAs acetate, propionate and butyrate) in plates as compared to tubes (**Figure S4C**).

The two butyrate-producing species *Agathobacter rectalis* and *Eubacterium ramulus* yielded significantly lower levels of formate in plate compared to tubes (9.5±2 vs. 12.5±1 mM, p≤0.05, and 1.1±0.03 vs. 3.5±0.1 mM, p≤0.001, respectively; **Figure S4C**). When grown in the chamber, the propionate-producing species *Phocaeicola dorei* showed higher conversion of succinate (1.8±0.03 vs. 3.0±0.1 mM, p≤0.001) to propionate (5.7±0.3 vs. 3.9±0.1 mM, p≤0.05; **Figure S4C**) compared to tubes. Collectively, these data highlight that the difference in atmosphere between the two cultivation techniques can impact the metabolic output during microbial cultivation. These differences were more pronounced in pure cultures than in complex communities. Specific procedures for testing fecal microbiota (**Supplementary file 1, Step 5.1**) or pure cultures (**Supplementary file 1, Step 5.2**) have been included in the high-throughput protocol.

### Formulating a growth medium to mimic the microbial composition of healthy adult feces in cultures

Next, we sought to formulate an optimal mix of C-sources for complementing bYCFA basal medium to maximize the compositional similarity between fecal cultures and feces from healthy human adults. By maintaining the main characteristics of the donor’s community in culture, we aimed to enhance the relevance of *in vitro* testing for evaluating the effect mediated by diet-, drug- and host-related factors. To this end, we cultivated fecal samples from eight healthy adults and examined whether the bacterial composition of feces could be maintained using complex mixtures of carbohydrates (3 g/L total; 3C and 6C) and mucin (+Muc), as compared to glucose and H_2_O (C-depleted control; **Figure 3A**). The composition of 3C was designed to mirror the C-sources present in the M2GSC medium (33% starch, 33% cellobiose and 33% glucose)^30^, while 6C contained the carbohydrates found in the Macfarlane medium (15% starch, 15% pectin, 15% xylan, 8% arabinogalactan, 8% guar gum and 38% inulin)^33^, a nutrient-rich medium that closely mimics the gut chyme of healthy adults, and is commonly used in continuous intestinal *in vitro* models.

**Figure 3:**
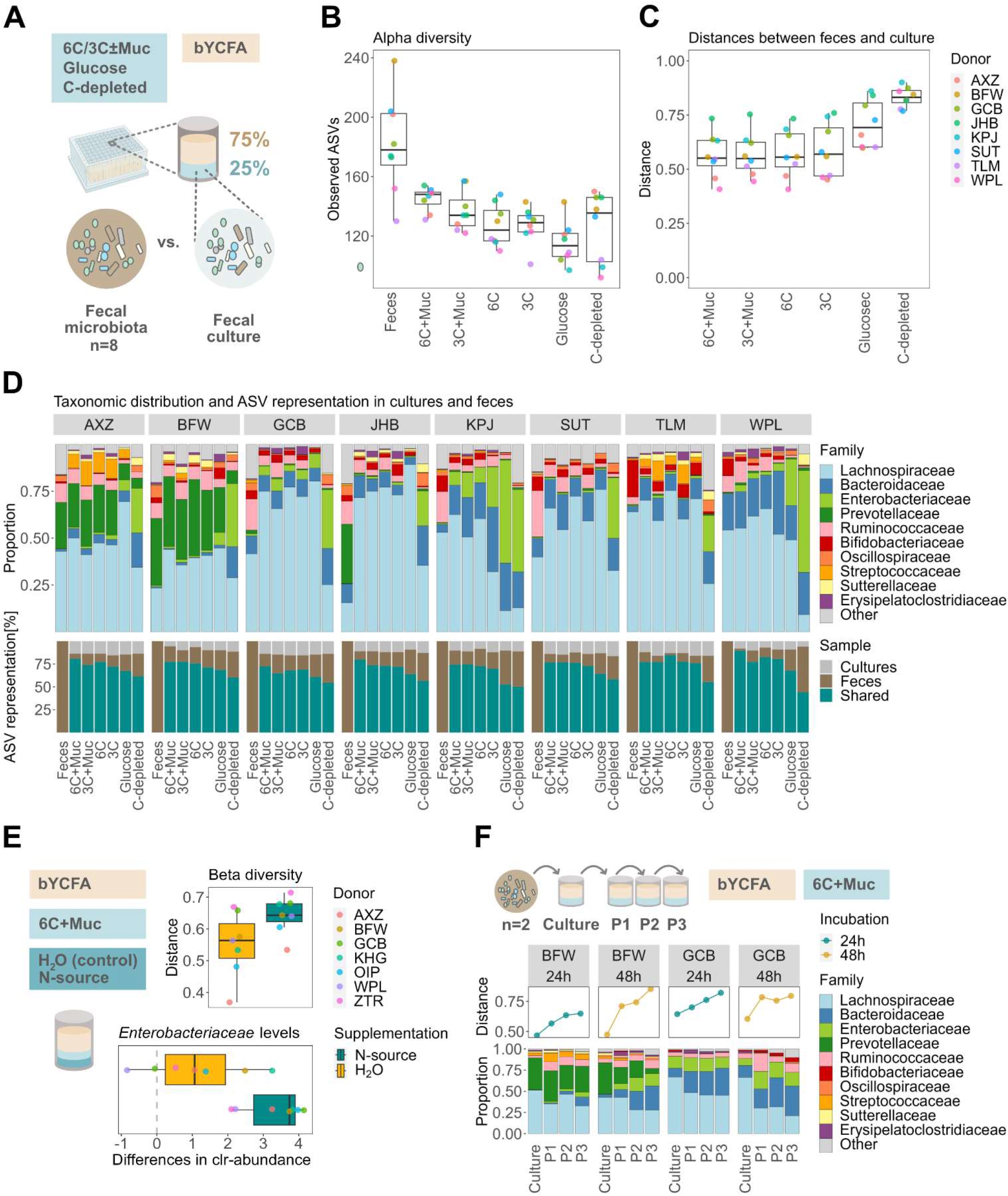
Formulation of a growth medium maintaining the microbial composition of healthy adult feces in cultures during high-throughput cultivation (48 h, 37°C) in bYCFA medium. **A)** Experimental setup for determining the optimal C-source mix to maintain the composition of the original feces in cultures. All analyses were performed after 48 h cultivation. **B)** Number of observed ASVs in feces and cultures with different C-sources. **C)** Bray-Curtis distances between the feces and the respective culture. **D)** Visual representation of the community composition (top) and the shared ASVs (with relative abundance >0.1%) between feces and cultures (bottom). **E)** Evaluation of the impact of additional N-source supplementation (amicase and yeast extract) on Bray-Curtis distances and on the difference in *Enterobacteriaceae* clr-abundance as compared to the feces. **F)** Bray-Curtis distances and composition of cultures re-inoculated (1%) into fresh bYCFA with 6C+Muc (24/48 h, 37°C, 3 passages, donors n=2). All analyses were performed on pooled samples from three independent replicates for each donor microbiota. A total of 11 distinct donors were used, *i.e.,* AXZ, BFW, GCB, JHB, KPJ, SUT, TLM, WPL, KHG, OIP and ZTR.

In terms of bacterial richness, the number of observed ASVs in feces (median of 178 among donors) was best maintained in culture conditions containing both mucin and complex C-sources, with the 6C+Muc condition exhibiting the highest number of ASVs (148), followed by 3C+Muc (134), 3C (129), 6C (124), and glucose (113.5) (**Figure 3B**). Though 135.5 ASVs were detected in the C-depleted control, a significant portion likely originated from resting cells from the fecal inoculum, as limited growth (OD_600_) was observed after 48 h (**Figure S5A**). When comparing the taxonomic composition of the fecal cultures to the original feces using Bray-Curtis distances (ranging from 0: all ASVs shared, to 1: no ASVs shared), the conditions containing complex C-sources demonstrated higher similarity to the feces (6C+Muc (0.55), 3C+Muc (0.55), 6C (0.56), and 3C (0.57)) than those supplemented with glucose (0.69) or H_2_O (0.83; **Figure 3C**). In line, shared ASVs (relative abundance >0.1%) between feces and cultures were highest in 6C+Muc (80±5%), followed by 3C+Muc (74±4%), 6C (73±5%), 3C (70±4%), and lowest with glucose (53±7%) or in the absence of a C-source (55±7%; **Figure 3D**). At the family level, the main fecal taxa were retained in cultures containing the complex C-sources (**Figure 3D**), except for donor JHB, for which a loss of *Prevotellaceae* abundance was observed after 48 h cultivation (0.6±0.3% among all complex C-source cultures vs. 32.2% in feces; **Figure 3D**). Notably, adding complex C-sources limited the relative abundance of *Enterobacteriaceae*, whose members have been associated with dysbiosis and were often reported to bloom during *in vitro* cultivation in the chamber^25,27,34^. The difference of *Enterobacteriaceae* levels between cultures and feces was smallest in the presence of 6C (clr-difference of 2.1, *i.e.*, 4.3-fold (2^2.1^) increase) and 6C+Muc (4.5-fold increase), and the largest in the presence of glucose (13.3-fold increase) and H_2_O (32.1-fold increase) (**Figure S5B**). Overall, among all cultures containing 6C+Muc, the maximum relative abundance of *Enterobacteriaceae* detected after 48 h was only 7.2% (Donor KPJ; **Figure 3D**).

When evaluating the metabolic outputs of fecal cultures, a significant increase in metabolite production was observed in the presence of C-sources (**Figure S5C**). Yet, substantial amounts of formate accumulated at the end of the cultivation (**Figure S5D**), indicating incomplete fermentation of the supplied C-source. We thus assessed whether the addition of N-sources could shift fermentation profiles towards end-metabolites^35^. We cultivated fecal samples from seven healthy adults in bYCFA with 6C+Muc and supplemented an additional 7.2 g/L amicase and 1 g/L yeast extract (**Figure 3E**).

With these additional N-sources, a reduction of formate levels after 48 h (from 14.4±3.8% to 1.3±2.6%) and a representative SCFA ratio^36^ was observed, with 61.5±5.1% acetate, 18.9±1.9% butyrate, and 16.4±5.4% propionate among all donors (**Figure S6A**). However, a higher N-source concentration did not help maintain the composition of the original feces. Though a minor effect was observed in terms of richness (123 ASVs with extra N-sources and 119 ASVs in control; **Figure S6B**), N-source addition increased Bray-Curtis distances between cultures and feces (extra N-source: 0.64; control: 0.56; **Figure 3E**). Similarly, the relative abundance of *Enterobacteriaceae* was enhanceded by supplementation with additional N-sources (**Figure 3E**). Therefore, we recommend complementing bYCFA with 6C+Muc without additional N-sources to better retain the compositional characteristics from healthy adult feces during cultivation (**Supplementary file 1**).

Lastly, we assessed whether communities cultured in bYCFA with 6C+Muc could be maintained through repeated passages to provide a setup for **studying**, *e.g.,* post-perturbation recovery patterns^25^. Two fecal cultures were re-inoculated (1% v/v) into fresh medium over three successive passages (P1, P2, P3) at two different intervals (24 or 48 h; **Figure 3F**). The successive passages increased the dissimilarity between the cultured communities and the original fecal samples (Bray-Curtis distances, **Figure 3F**). Despite a similar overall divergence of community structure between 24 and 48 h intervals, there was a notable difference in taxonomic composition and metabolic activity. A shorter incubation time of 24 h compared to 48 h resulted in an increase in the clr-abundance of *Enterobacteriaceae* between initial cultures and subsequent passages (**Figure S7A**) and mitigated the decline of *Prevotellaceae* in both donors (**Figure S7A**). Conversely, higher levels of intermediate metabolites (*i.e.,* formate, succinate) were detected in 24 h compared to 48 h passages (**Figure S7B**). Although successive re-inoculation of fecal cultures induced shifts in community structure after each passage, this approach still shows promise for studying recovery dynamics in a high-throughput manner.

### Treating fecal cultures with dietary fibers or drugs results in physiologically relevant microbial responses

Finally, we aimed to validate our *in vitro* high-throughput cultivation protocol by testing whether supplementation with specific dietary fibers (resistant dextrin and soluble starch; **Figure 4A**) or drugs (omeprazole, ciprofloxacin and 5-fluorouracil (5-FU); **Figure 4D**) could induce microbial changes that are consistent with previous *in vivo* findings.

**Figure 4:**
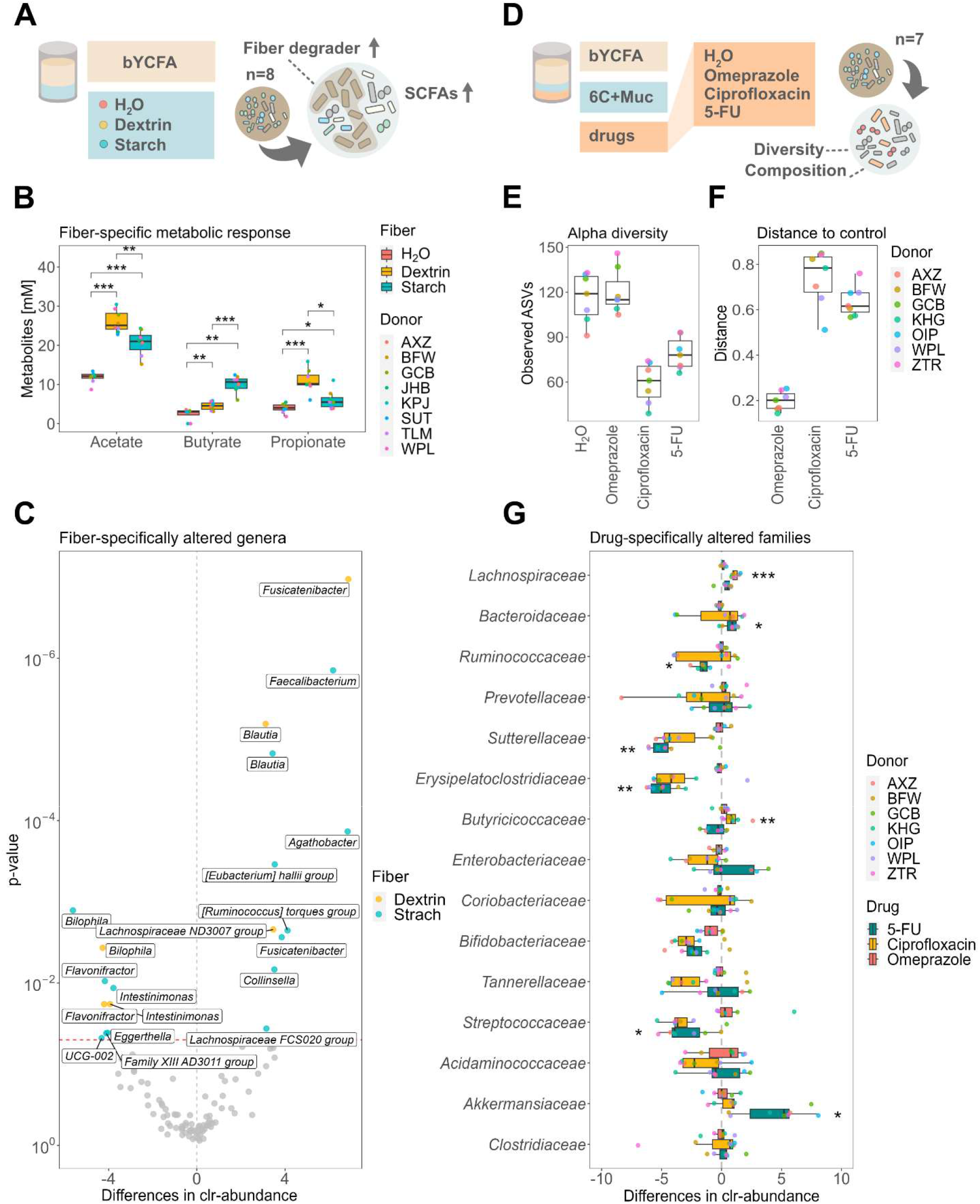
Treatment with dietary fibers (donor n=8) or drugs (donor n=7) elicits physiologically relevant changes in the fecal cultures’ metabolism and community composition (bYCFA, 48 h, 37°C). A) Experimental setup to test the fiber-specific response of fecal cultures. All analyses were performed after 48 h cultivation. B) SCFA production by fecal cultures when bYCFA was supplemented with resistant dextrin (3 g/L), soluble starch (3 g/L) or no C-source (H_2_O as control). C) Median clr-differences of taxa significantly differed between each fiber (dextrin, starch) and H_2_O. Genera significantly different (p≤0.05 and clr-difference >3) are labeled and located above the red line. D) Experimental setup to test the effect of drugs (*i.e.,* omeprazole, ciprofloxacin and 5-FU) as compared to the control (H_2_O) in bYCFA supplemented with 6C+Muc. All analyses were performed after 48 h cultivation. E) Number of observed ASVs. F) Bray-Curtis distances between controls and the drug-treated cultures. G) Clr-differences of the 15 most abundant families. Significantly altered families, as compared to the control, are marked by stars. All the analyses were performed on pooled samples from incubations for each donor microbiota. Statistically significant results are marked by stars, with * indicating p≤0.05, ** p≤0.01 and *** p≤0.001.

We chose resistant dextrin (3 g/L) and soluble starch (3 g/L) as dietary fibers due to their well-known ability to modulate the gut microbiota and mediate propiogenic^37^ and butyrogenic^38–40^ effects, respectively. Using feces from eight healthy donors, we performed enrichment experiments in bYCFA containing each fiber as the sole C-source. Among all donors, we found significantly higher levels of propionate after 48 h in the presence of dextrin (10.9±2.9 mM) compared to starch (6.0±2.4 mM; p≤0.05) or the control without C-sources (3.9±1.2 mM; p≤0.001). Conversely, starch supplementation led to significantly higher levels of butyrate (10.1±2.1 mM) compared to dextrin (4.5±1.1 mM; p≤0.001) or the control (2.4±1.5 mM; p≤0.01) (**Figure 4B**). At the taxonomic level, resistant dextrin specifically increased the abundance of *Fusicatenibacter* (p≤0.001) and *Lachnospiraceae* ND3007 group (p≤0.01) among all donors (**Figure 4C**), consistent with previous *in vitro* and *in vivo* studies^24,41^. Soluble starch supplementation enriched *Faecalibacterium* (p≤0.001) and *Agathobacter* (p≤0.001; formerly *Eubacterium*; **Figure 4C**) as shown in previous human trials^17,39,40,42^. *Blautia* was enriched by both fibers (p≤0.001; **Figure 4C**).

Many commonly-used medications have been shown to disrupt gut microbial communities^6,43^. Reciprocally, microbial activities can alter drug efficacy and toxicity^44–46^. To test the direct interaction between the gut microbiota and drugs, we exposed the feces from seven healthy donors (in bYCFA with 6C+Muc) to a single dose of 11 μM omeprazole (proton-pump inhibitor), 50 μM ciprofloxacin (antibiotic) or 50 μM 5-FU (cytostatic), at their expected intestinal concentrations^25,47,48^.

Compared to the control cultures, treatment with ciprofloxacin and 5-FU negatively impacted the microbial communities’ diversity (**Figure 4E**) and structure (**Figure 4F**). Ciprofloxacin led to a relative increase in *Butyricicoccaceae* (p≤0.01) and *Lachnospiraceae* (p≤0.001) (**Figure 4G**). This reduction in community diversity and persistence of *Lachnospiraceae* agrees with previous findings in mice^25^ and ciprofloxacin-treated patients^49^. Treatment with 5-FU inhibited *Sutterellaceae* (p≤0.01) and *Erysipelatoclostridiaceae* (p≤0.01) and promoted the relative abundance of *Akkermansiaceae* (p≤0.05) (**Figure 4G**). The reduction in diversity and the drastic bloom of *Akkermansiaceae* in response to 5-FU have been demonstrated in several mice studies^50–52^. Conversely, we observed no major effect of omeprazole on fecal cultures at the tested concentration (**Figures 4E and 4F**). This common proton pump inhibitor has been shown to decrease the richness, enrich the abundance of oral bacteria (e.g., *Streptococcaceae*) in the gut, and increase the risk of enteric infections *in vivo*^53^. The effect of omeprazole on the gut microbiota was proposed to be primarily mediated by the increase in gastric pH, which disrupts the barrier between the upper and lower gastrointestinal tract, rather than through a direct effect of the molecule itself^53^. Our data align with this hypothesis, with no direct impact observed on the microbiota. It is worth noting that omeprazole was dissolved in DMSO at a final concentration of 0.2% DMSO in fecal culture. At this concentration, DMSO did not affect the community diversity and structure (**Figure S8**), indicating that it can serve as an optimal solvent for testing compounds with low solubility in an aqueous phase.

## DISCUSSION

Developing a well-controlled protocol for cultivating stool-derived microbial communities can help advance our understanding of the causal interactions between gut microbes, diet, drugs, and host physiology^54^. However, the throughput of current anaerobic cultivation techniques is limited and insufficient to capture the complex nature and individuality of the gut microbiota, with heterogenous compositions among individuals^16–19^, variations in activities at the strain level^20–23^, and the influence of environmental factors on ecological and physiological dynamics^6,10^. A few pioneering studies have reported development of anaerobic high-throughput systems for cultivating stool-derived gut microbiota to generate personalized culture collections of gut microbes^55,56^, map the ability of the human gut microbiota to metabolize small molecule drugs^27^ or evaluate the response of fecal communities to specific stimuli^25,26^. These studies have highlighted the promise of high-throughput *in vitro* modeling by demonstrating a strong correlation between *in vitro* and *in vivo* gut microbiota responses^25,26^. Here, we addressed practical, technical, and biological gaps in fecal cultivation that we consider essential to create a widely applicable, reproducible and accessible protocol to advance the field of gut microbiota research.

First, we developed a protocol that relies solely on commonly-available laboratory materials and equipment, making it easily implementable in various research settings. The core requirement is an anaerobic chamber equipped with a pH meter, an incubator, and multichannel pipettes. We have selected 96-deepwell plates for cultivation, as they offer various advantages; they are reusable and provide a high throughput capacity (60 wells available per plate for testing) with a sufficient volume of samples for analyses (2 mL). Further, the small surface area of wells minimizes evaporation and gas transfer rates, thus reducing the effect of potential O_2_ contamination^57^. The main limitation of 96-deepwell plates is their inability to support online monitoring of absorbance or fluorescence using plate readers.

Second, we comprehensively characterized confounding factors associated with anaerobic chamber cultivation. We showed that the gas phase of the chamber i) might contain residual O_2_, ii) alters the pH of the medium, and iii) modulates the C-metabolism of intestinal microbes. To prevent oxygen contamination, we recommend properly preparing the chamber by monitoring H_2_ levels and regenerating the palladium catalysts. Our protocol also includes a step of boiling and flushing (CO_2_ or N_2_) of the basal medium prior to sterilization (**Supplementary file 1**), similar to what is done with the Hungate technique^14^. While it is theoretically possible to skip these steps and generate anaerobiosis by pre-reducing a sterile aerobic medium inside the chamber for several days before the experiments^25,27^, we observed a limited bloom of facultative aerobes, such as *Enterobacteriaceae*, with our protocol compared to other high-throughput cultivation studies^25,27^. We also highlighted the negative impact of a weak reducing atmosphere on the growth of intestinal microbes (**Figure 1B**). Therefore, we strongly recommend boiling and flushing the basal medium before sterilizing to ensure strict anaerobiosis. Regarding pH, we highlight that the equilibrium between gaseous and dissolved CO_2_ and the concentration of NaHCO_3_ in the solution influenced the pH in the chamber. Thus, the bicarbonate buffer is more challenging to control in an open system, such as 96-deepwell plates where CO_2_ can escape. With a buffering range between pH 5.1 to 7.1^32^, it can maintain the pH in the range of the human adult colon, from 5.7 (proximal) to 6.7 (distal)^58^, with colonic pH values as low as 5.0 in some healthy individuals^59^. In addition, the bYCFA basal medium contains a phosphate buffer to maintain the pH between 6.4 and 7.4. Concerning the impact of the gas phase on microbial metabolism, we highlighted a few biological differences between the plate-based anaerobic chamber and the classic tube-based Hungate technique. The typically lower CO_2_ concentration in the chamber (∼10%) compared to the initial concentration in tubes (∼100%) mediated an increase of succinate to propionate conversion in pure cultures of *P. dorei* (**Figure S4C**)^60,61^. Moreover, the plate-based system allows microbially-produced gases to be released, including H_2_, a common growth-limiting factor that accumulates in tube cultures during the fermentation process^62^ and promotes accumulation of intermediate metabolites^63^. This study supports this finding, as tube cultures exhibited a higher accumulation of intermediate products (*i.e.,* formate) than plate cultivation (**Figures 2C**, **S3C and S4C**). Overall, the typical gas composition in the anaerobic chamber (*i.e.,* 5% H_2_, 10% CO_2_, and 85% N_2_) is close to the physiological conditions reported *in vivo* for H_2_ and CO_2_, with concentrations of 2.9±0.7% and 9.9±1.6%, respectively^64^.

Third, we designed a reproducible and modular medium. The basal medium bYCFA was formulated to be heat-stable (sterilization via autoclave) and to contain as few undefined ingredients as possible to increase batch-to-batch reproducibility. This basal medium can be flexibly complemented with C-sources or any other molecules to test, and the pH can be easily adjusted, allowing for testing of a wide range of culture conditions with minimal medium preparation effort. Therefore, the effects of a combination of factors can be easily tested and incorporated in the experimental design (**Supplementary file 1**). Here, we showcased two examples of experiments. We first complemented bYCFA with defined fibers to identify taxa associated with functional niches (**Figures 4A, 4B and 4C**). This approach corroborated results from a previous tube-based enrichment study^24^ while allowing evaluation of more donors and conditions simultaneously. We then maintained fecal culture communities resembling the original feces by complementing bYCFA with an optimized mix of C-sources (6C+Muc; **Figure 3B, 3C and 3D**), and tested microbial responses to antibiotic and non-antibiotic drugs (**Figures 4D, 4E, 4F and 4G**). This approach can be extended to study the effects of other xenobiotics *(e.g.,* pollutants), dietary compounds (*e.g*., N-sources, vitamins), or host-derived factors (*e.g*., pH, redox potential, oxidative stress, bile salts, antimicrobial peptides). Note that molecules insoluble in water can be dissolved in DMSO, with no effect on the microbial community at a concentration of 0.2% (**Figure S8**) and potentially up to 1%^65^.

Finally, we integrated all technical procedures and practical solutions into a simple yet comprehensive step-by-step protocol (**Supplementary file 1**). This protocol provides a high level of detail and encompasses key features like flexibility (*e.g*., testing of various molecules and conditions alone or in combination; complex communities or pure cultures), reproducibility (*e.g*., reduced media complexity; preparation of stable stock solutions), and simplicity (*e.g*., minimized hands-on time; easy pH adjustment; reduced evaporation rate). In conclusion, our high-throughput cultivation protocol is a valuable tool for fundamental and applied gut microbiota research. It enables rapid and reproducible characterization of diet-, host-, or drug-specific effects, microbe-microbe interactions^66^, strain-specific^20,23^ and donor-specific^19^ responses and activities. The protocol can also be applied to cost-efficient screening of drug metabolism by the gut microbiota^46,47^, pre-selection of treatments or dosage prior to costly animal or human trials^67^, development of personalized microbiota-targeted nutritional interventions^68^, identification of responders and non-responders to drugs and prebiotics^19^ and for discovery and investigation of novel live biotherapeutics^69^.

## ACKNOWLEDGMENTS

This work was supported by a grant from the Swiss Innovation Agency (51128.1 IP-LS) in collaboration with PharmaBiome AG. This work was also supported by an ETH Research Grant ETH-38 20-1. We thank Alfonso Die (ETH Zürich) for their assistance with HPLC-RI analyses. We thank Florentin Constancias for his support with statistical analysis. We thank Anna Greppi for her valuable feedback during the preparation of the manuscript. We thank Roquette for providing NUTRIOSE® (resistant dextrin). Data were generated in collaboration with the Genetic Diversity Centre (GDC), ETH Zurich. Graphic elements BioRender.com. have been used to create Figures 1,2 and 3.

## AUTHOR CONTRIBUTIONS

BP, CL, TW, and GL conceptualized the project and provided financial support. JZ, BP and GL planned the experiments. JZ, SP, DM, CM performed wet-lab experiments and analyses. JZ, PB, GL curated the data. JZ and PB performed data analysis. JZ, BP, GL, CL interpreted the data. JZ, SP and BP wrote the step-by-step protocol. JZ and BP wrote the manuscript. All authors critically reviewed the manuscript. All authors contributed to the article and approved the submitted version.

## INCLUSION AND DIVERSITY

We support inclusive, diverse, and equitable conduct of research.

## METHODS

### Experimental model and subject details

#### Human fecal samples

14 healthy donors (7 females, 7 males, age ranging between 26 and 36) consented to provide one or more fresh fecal samples. Donors were excluded if they had taken any antibiotics in the past six months. Donor identity was ensured by using randomly generated three-letter codes. The ethics committee of ETH Zurich exempted this study from review as the sample collection was processed in an anonymized and non-interventional manner.

#### Bacterial strains

A panel of ten human intestinal strains, representing a panel of fiber and intermediate metabolite utilizers, was acquired from the German Collection of Microorganisms and Cell Culture GmbH (DSMZ, Braunschweig, Germany) and the strain collection of PharmaBiome AG (Zürich, Switzerland; **Table S2**). Strains were stored at −80°C in anaerobic glycerol stocks (25%, v/v) and reactivated by inoculating (1.25%, v/v) Hungate tubes containing bYCFA (pH 6.8) supplemented with the respective carbon sources (final concentrations of 3 g/L for fibers, 30 mM for glucose and intermediate metabolites; **Table S2**). Pre-cultures were incubated for 24 h at 37°C before experiments.

### Method details

#### Growth media and anaerobic procedures

To cultivate fecal microbiota and pure cultures, we modified the yeast casitone fatty acid medium (YCFA)^31^ and the M2GSC^30^ medium. Modifications included a tenfold reduction of N-sources, as previously reported^24^. The resulting basal minimal media, referred to as basal YCFA (bYCFA) and modified M2GSC (mM2), consisted of (L^−1^): 1 g amicase (Sigma-Aldrich Chemie GmbH, Buchs, Switzerland), 1.25 g yeast extract (Lesaffre, Marcq-en-Barœul, France), 0.5 g meat extract (Sigma-Aldrich), 150 mL mineral solution I (0.52 g K_2_HPO_4_), 150 mL mineral solution II (0.52 g KH_2_PO_4_, 0.9 g NaCl, 0.9 g (NH_4_)_2_SO_4_, 90 mg MgSO_4_, 90 mg CaCl_2_), 1 g L-cysteine HCl, and 4 g NaHCO_3_. The medium mM2 further contained 300 mL rumen fluid (centrifuged and stored at −20°C after collection; Vetsuisse Faculty University of Zurich, Zürich, Switzerland). The rumen fluid was substituted in bYCFA with defined compounds, *i.e.,* 0.2 mL hemin stock solution (10 μg hemin from bovine, Sigma-Aldrich), 0.1 mL vitamin solution (10 μg biotin, 10 μg cobalamin, 30 μg 4-aminobenzoic acid, 50 μg folic acid and 150 μg pyridoxamine, Sigma-Aldrich) and 6.2 mL of volatile fatty acids solution (2.11 mL acetic acid (Sigma-Aldrich), 0.74 mL propionic acid (Sigma-Aldrich), 0.12 mL valeric acid (VWR International AG, Dietikon, Switzerland), 0.12 mL isovaleric acid (Sigma-Aldrich), and 0.12 mL isobutyric acid (Sigma-Aldrich) in 3.1 mL 5 M NaOH). Since the basal medium was complemented with (heat-stable/-sensitive) supplements after sterilization (**Figure 1A**), it was thus prepared 1.33-fold concentrated (**Table S3**). All compounds except L-cysteine.HCl and NaHCO_3_ were dissolved in 75% of the final volume (**Table S3**). The basal medium was adjusted to pH 7, boiled (10 min), and the remaining compounds were added after cooling to approximately 60°C while constantly flushing with CO_2_ (or N_2_). After another 10 min, the medium was transferred into DURAN® Pressure flasks (Huberlab AG, Aesch, Switzerland) or Hungate tubes (Milian, Vernier, Switzerland) and autoclaved.

To complement the basal medium, a range of heat-stable supplement solutions were prepared (**Table S3**). For fecal microbiota cultivation, we supplemented the media with different concentrated (4-fold or 8-fold; **Table 3**) C-source solutions resulting in a final concentration of total 3 g/L of the C-sources when combined with the (1.33-fold) concentrated basal medium. C-source solutions included resistant dextrin (NUTRIOSE® FB06; Roquette, Lestrem, France), starch (soluble starch from potato; Sigma-Aldrich), glucose (Sigma-Aldrich), a mix of six sugars (referred to as 6C; 0.45 g/L starch, 0.45 g/L pectin (pectin from citrus peel; Sigma-Aldrich), 0.45 g/L xylan (xylan from oat spelt; Angene, London, United Kingdom), 0.24 g/L arabinogalactan (from larch wood; Sigma-Aldrich), 0.24 g/L guar (Sigma-Aldrich) and 1.14 g/L inulin (Orafti GR; Beneo, Mannheim, Germany) or a mix of three sugars (referred to as 3C; 1 g/L starch, 1 g/L cellobiose (Sigma-Aldrich) and 1 g/L glucose). The 6C and 3C mixes were also tested with mucin from porcine stomach type II (0.3 g/L final concentration; Sigma-Aldrich). Heat-stable carbohydrates were also supplemented to basal bYCFA for cultivating pure bacterial strains, as described in **Table S2**. Additional heat-stable supplements included (8-fold) concentrated N-source solutions, which resulted in a final concentration of 8.2 g/L amicase and 2.25 g/L yeast extract. Heat-stable compounds were dissolved in dH_2_O, adjusted to pH 7, boiled for 10 min, cooled under N_2_ and autoclaved separately soon after.

In addition, several heat-sensitive compounds were tested (**Table S3**), including antibiotic and non-antibiotic drugs at final concentrations of 50 μM ciprofloxacin (Sigma-Aldrich), 11 μM omeprazole (Sigma-Aldrich) and 50 μM 5-fluorouracil (Sigma-Aldrich). Concentrated drugs (8-fold) were dissolved in anaerobic dH_2_O (except for omeprazole which was dissolved in dimethyl sulfoxide (DMSO); 0.2% final concentration) in the anaerobic chamber, adjusted to pH 7 and filter sterilized.

The day before the experiment, the autoclaved basal medium was equilibrated with the gas phase of the anaerobic chamber (10% CO_2_, 5% H_2_, and 85% N_2_, Coy Laboratory Products Inc., Grass Lake, MI, United States) under constant stirring, without the lid but covered with Breathe-Easier Sealing film (Diversified Biotech, Dedham, MA, United States). On the day of the experiment, the pH was adjusted to 6.8 (if not indicated otherwise) by adding HCl 3 M based on a pre-determined titration curve (**Figure 1E**) and was mixed with the corresponding heat-stable/sensitive supplements (**Table S3**).

A complete step-by-step description of the preparation of the basal bYCFA medium and heat-stable/sensitive supplement solutions for the high-throughput cultivation in 96-deepwell plates (Milian, Vernier, Switzerland) is presented in **Supplementary file 1**. For gas-tight tube-based culture using the Hungate technique, anaerobiosis was maintained by applying a strict anaerobic technique as described previously^70^. Syringes and needles were pre-flushed with filter-sterilized CO_2_ before being used to inoculate carbon source solutions or bacteria.

#### Fecal sample inoculation, cultivation and sampling procedure

Healthy adult donors provided a fresh fecal sample in a plastic container containing an Oxoid™ AnaeroGen™ pouch (Thermo Fisher Diagnostics AG, Pratteln, Switzerland) to generate anaerobiosis during transport to the laboratory. Within 3 hours, the samples were transferred into the anaerobic chamber. Approximately 1 g of fecal matter was transferred using a plastic spoon into a 50 mL Falcon tube and homogenized with 9 mL of anaerobic phosphate buffer (pH 7.0). The resulting 10^−1^ dilution was then further diluted to 10^−4^ and was inoculated (1%, v/v) into a plate or tube containing previously pH-adjusted basal medium (pH 6.8) complemented with respective supplements (**Supplementary file 1, Step 5.2**). Unless otherwise stated, working volumes of 2 mL and 8 mL of medium were used in 96-deepwell plates and gas-tight Hungate tubes, respectively. After incubation for 48 h at 37°C, cultures were transferred out of the chamber and centrifuged (4’700 rpm, 20 min and 4°C). Cell pellets and supernatants were recovered and stored separately (−80°C) until further processing (**Supplementary file 1, Step 8**).

#### Anaerobic chamber preparation and its impact on bacterial growth

We assessed the effects of anaerobic chamber preparation on the growth of gut microbes in pure cultures. Growth kinetics of *Bacteroides thetaiotaomicron* DSM 2079 and *Faecalibacterium prausnitzii* DSM 17677 were monitored in an anaerobic chamber that was handled “strictly” or “loosely”. The “strictly” handled chamber (**Supplementary file 1**, **Step 2**) consisted of regenerated palladium catalysts and a gas atmosphere replaced with fresh gas mix (10% CO_2_, 5% H_2_, and 85% N_2_), resulting in 0-20 ppm O_2_ and >2.5% H_2_ (monitored with an Anaerobic Monitor CAM-12, Coy Laboratory Products Inc.). To mimick a “loosely” handled chamber, non-regenerated palladium catalysts were introduced and gas mix was replaced with N_2_ until the H_2_ level was less than 1.5%.

Cultures (adjusted to OD_600_ ∼0.2) were inoculated (1% v/v) into 200 µL bYCFA (30 mM glucose, pH 6.5) in a 96-well plate (SPL Life Sciences Co. Ltd., Gyeonggi-do, South Korea). To demonstrate the effect of chamber handling, the incubation (36 h, 37°C) and monitoring of growth kinetics (Tecan Infinite M200 PRO plate reader; Tecan Group Ltd., Männedorf, Switzerland) was performed in a “strictly” or “loosely” handled anaerobic chamber. Each condition was tested in three biological replicates. The maximum specific growth rate (μ_max_) was calculated according to first-order growth kinetics.

#### pH dynamics and bicarbonate buffering system in the anaerobic chamber

We characterized the relationship between pH, NaHCO_3_ concentration, and CO_2_ levels (0% or 100% in gas-tight tubes, compared to 10% in the anaerobic chamber) by measuring the pH of the basal medium containing different NaHCO_3_ concentrations (0.4, 1, 2, 4 g/L). When required, pH dynamics were monitored over time. Measurements were performed using a pH microelectrode (diameter 6 mm; Metrohm AG, Herisau, Switzerland) and a pH meter (type 913; Metrohm AG). pH was measured in a 96-deepwell plate filled with 1.5 mL basal medium. A titration of bYCFA medium with HCl 3 M was performed and served as a calibration to support the pH adjustment of bYCFA in the chamber (**Supplementary file 1**, **Step 6**).

#### Comparison of the cultivation techniques (96-deepwell plates vs. gas-tight tubes) using fecal cultures and pure bacterial cultures

Two stool-derived gut microbiota (donors ABX and BCY) and eight intestinal strains (**Table S2**) were assessed for their growth and metabolic capabilities using plate- and tube-based techniques. Plates and tubes were filled with bYCFA or mM2 medium, complemented with the corresponding C-sources (**Table S3**), and inoculated (1% v/v) with fecal dilutions or pre-cultures of pure strains. After incubation (48 h, 37°C), 200 µL of cultures were transferred into a 96-well plate for OD_600_ measurement. Supernatants and the cell pellets were recovered (4’700 rpm, 20 min and 4°C) for organic acids quantification and 16S rRNA sequencing analysis, respectively.

#### Optimal medium composition to culture healthy adult feces and the response of individuals’ gut microbiota to fibers and drugs during high-throughput cultivation

We inoculated fecal dilutions from eight (C-source evaluation) or seven (N-source evaluation; fibers and drugs testing) adult donors in 96-deepwell plates containing bYCFA complemented with the corresponding supplements listed in **Table S3**.

Previous experiments (Validation of 96-deepwell plates with gas-tight tubes) resulted in small compositional differences within replicates (median Bray-Curtis distance 0.11) compared to the between-sample difference (median Bray-Curtis distance 0.72; **Figure S9**). Thus, given the low compositional variability, replicates were pooled by transferring 0.5 mL from each culture into a fresh plate (**Supplementary file 1**, **Step 8)** to reduce sample analysis time, cost and storage space. Two pools were prepared, one of which was stored immediately at −80°C, while the other was processed as described above.

#### DNA extraction and 16S rRNA sequencing

Two different extraction and sequencing protocols were applied. Specifically, for the initial comparison of the cultivation techniques (96-deepwell plates vs. gas-tight tubes), the following 16S rRNA sequencing pipeline was used: DNA extraction of fecal cultures was performed using the Maxwell® RSC instrument and the Maxwell® RSC PureFood GMO and Authentication Kit (Promega AG, Dübendorf, Switzerland). Library preparation, amplification, and paired-end (2x 300) sequencing of the hyper-variable regions V3/V4 of the genomic 16S rRNA gene with the primers (341F: 5“-CCTACGGGNBGCASCAG-3”; 806bR: 5“-GGACTACNVGGGTWTCTAAT-3”; Integrated DNA Technologies, Leuven, Belgium) was performed by StarSeq GmbH (Mainz, Germany) using Illumina MiSeq. Cell pellets originating from later experiments were sequenced as described below.

Cell pellets resulting from later experiments (optimization of medium composition and fiber- and drug-testing) were pre-processed using the Maxwell® HT 96 gDNA Blood Isolation Kit (Promega AG, Dübendorf, Switzerland) combined with a FastPrep homogenizer (FastPrep-24TM; MP Biomedicals, Illkirch-Graffenstaden, France), and followed by DNA purification using KingFisher Flex instrument (Thermo Fisher Scientific, Waltham, MA, United States), according to the manufacturers’ instructions. Amplification and barcoding of the V4 hypervariable region of the 16S rRNA gene was performed using a PCR with the barcoded primer 515F (5′-GTGCCAGCMGCCGCGGTAA-3′) and 806R (5′-GGACTACHVGGGTWTCTAAT-3′)^71,72^. The pooled library was then sequenced using the amplifying primers (515F and 806R) and the Illumina MiSeq platform^73^ at the Genetic Diversity Centre (ETH Zurich, Switzerland).

### Quantification and statistical analysis

#### Organic acids quantification

Short-chain fatty acids (SCFA; acetate, propionate and butyrate) and intermediate metabolites (formate, lactate and succinate) were quantified by high-performance liquid chromatography equipped with a refractive index detector (HPLC-RI). Briefly, approximately 300 µL supernatant was passed through a 0.2 µm nylon membrane filter (AcroPrep™ Filter plate; Pall Corporation, New York, United States; or syringe filters; Merck & Cie KmG, Schaffhausen, Switzerland). Analysis with a sample volume of 40 µL was carried out using a Chromaster HPLC-System (Hitachi, Tokyo, Japan) equipped with a SecurityGuard Cartridge Carbo-H (4×3.0 mm; Phenomenex Inc., Torrance, CA, United States) and a Rezex ROA-Organic Acid H+ (300×7.8 mm; Phenomenex Inc.) column. Elution was performed at 80°C under isocratic conditions (10 mM H_2_SO_4_, flow rate 0.6 mL/min), and analytes were quantified using a refractive index detector Chromaster 5450 (Hitachi). Data were processed using the Chromaster software (Hitachi).

All listed compound concentrations were corrected with blank values of the basal medium to estimate the net production of SCFA and intermediates. To quantify total metabolites, the concentration of the medium-corrected concentrations of measured metabolites were summed. Metabolite fractions were calculated by dividing the medium-corrected concentration of the respective metabolite by total metabolites.

#### Data analysis and visualization

Data were analyzed using R software (v4.2.0) and visualized using the ggplot package (3.3.6). Correlation lines were plotted using ggpmisc (0.5.1). Data were evaluated for normality using the Shapiro-Wilk Normality Test and when p>0.05, the comparison of means was performed using a t-test. Otherwise, a Wilcoxon test using the rstatix package (0.7.0) was used.

### 16S rRNA sequencing analysis

Sequencing reads were processed using the metabaRpipe R package (v0.9)^74^. In brief, reads were inferred with the DADA2 R package (v1.14.1)^75^ with read filtering set to c(260, 250) and maximal error rates set to c(3,4). Bimeras were then removed and taxonomy was assigned to ASVs using the SILVA database (v138.1)^76,77^. The communities were then analyzed using the phyloseq package (v1.40.0) of the R software (v4.2.0)^78^. Alpha and beta-diversity analyses were done on rarefied samples (with a minimal rarefication depth of 1556 reads). Data meeting the criteria of homoscedasticity were evaluated with a permutational multivariate analysis using the adonis2 function of the vegan package (v2.6-4). Differential abundance analysis was performed on non-rarified data using ALDEx2^79^. Using the aldex.clr function from the mia package (v1.5.17), the data was centered log(2)-ratio (clr) transformed using 1000 Dirichlet Monte-Carlo (DMC) instances. Significantly altered taxa were identified based on the p-value of Welch’s t-test.

## SUPPLEMENTARY FIGURES

**Figure S1:**
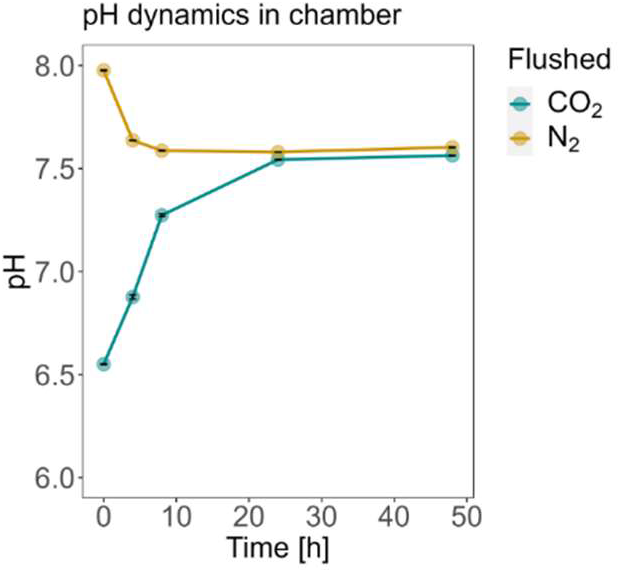
pH dynamics of CO_2_- and N_2_-flushed mM2 in 96-deepwell plate during equilibration in the anaerobic chamber (uninoculated, n=3).

**Figure S2:**
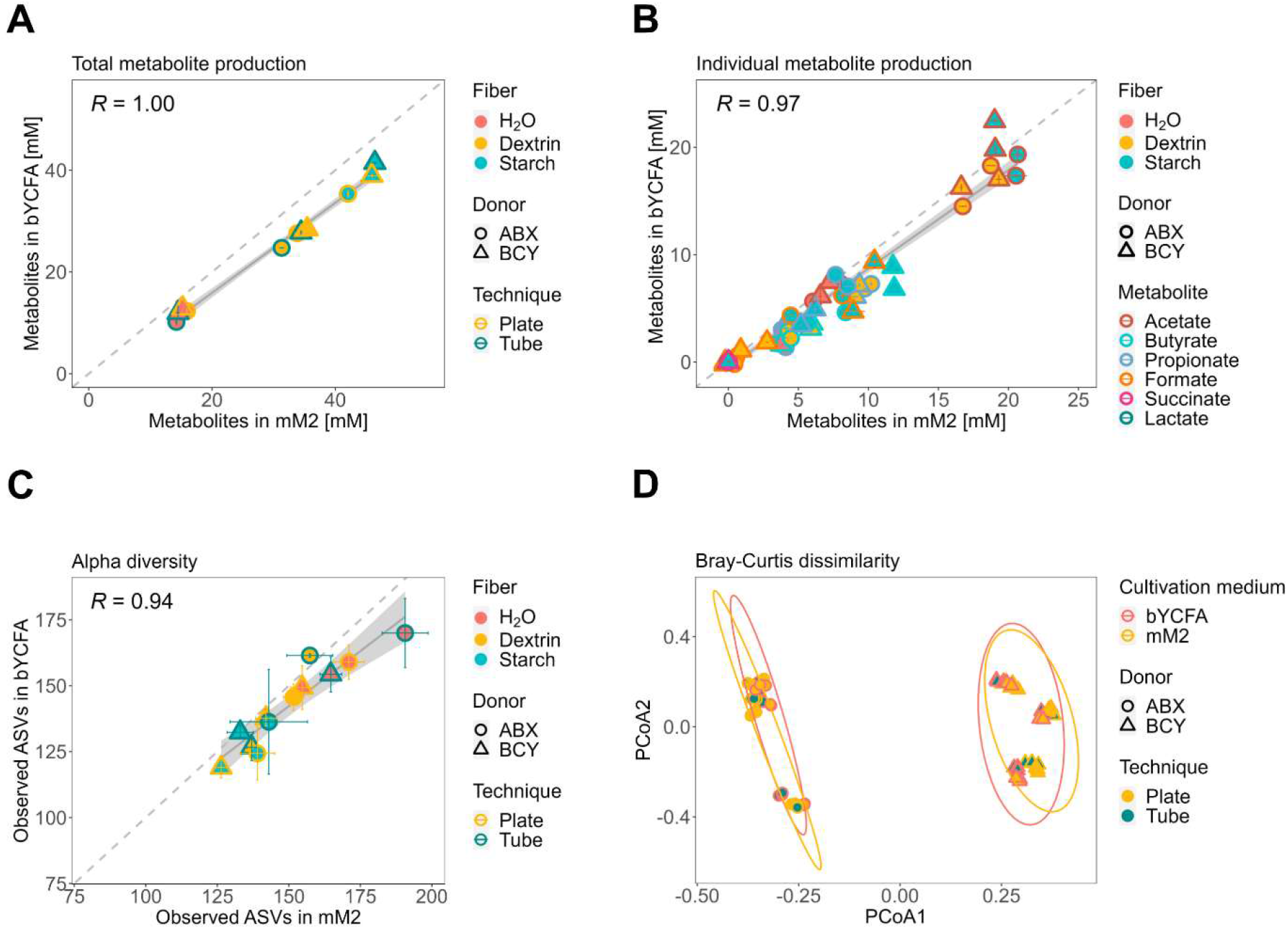
Impact of basal media (bYCFA vs. mM2) on metabolism and community structure of fecal-derived gut microbiota (donor ABX and BCY) cultivated 96-deepwell plates and gas-tight tubes for 48 h, 37°C, in the presence of resistant dextrin (3 g/L), soluble starch (3 g/L) or H_2_O as control (technical triplicates). All analyses were performed after 48 h cultivation. **A)** Correlation of the total metabolite production. Total metabolites were calculated by summing the medium-corrected concentrations of organic acids (SCFA and intermediates). **B)** Correlation of individual metabolites. Points represent means of blank corrected concentrations and bars represent standard deviations. **C)** Correlation of observed ASVs. Points represent means and bars represent standard deviations. **D)** PCoA plot of Bray-Curtis distances with 95% confidence ellipses (calculated separately for each donor) for the two basal media.

**Figure S3:**
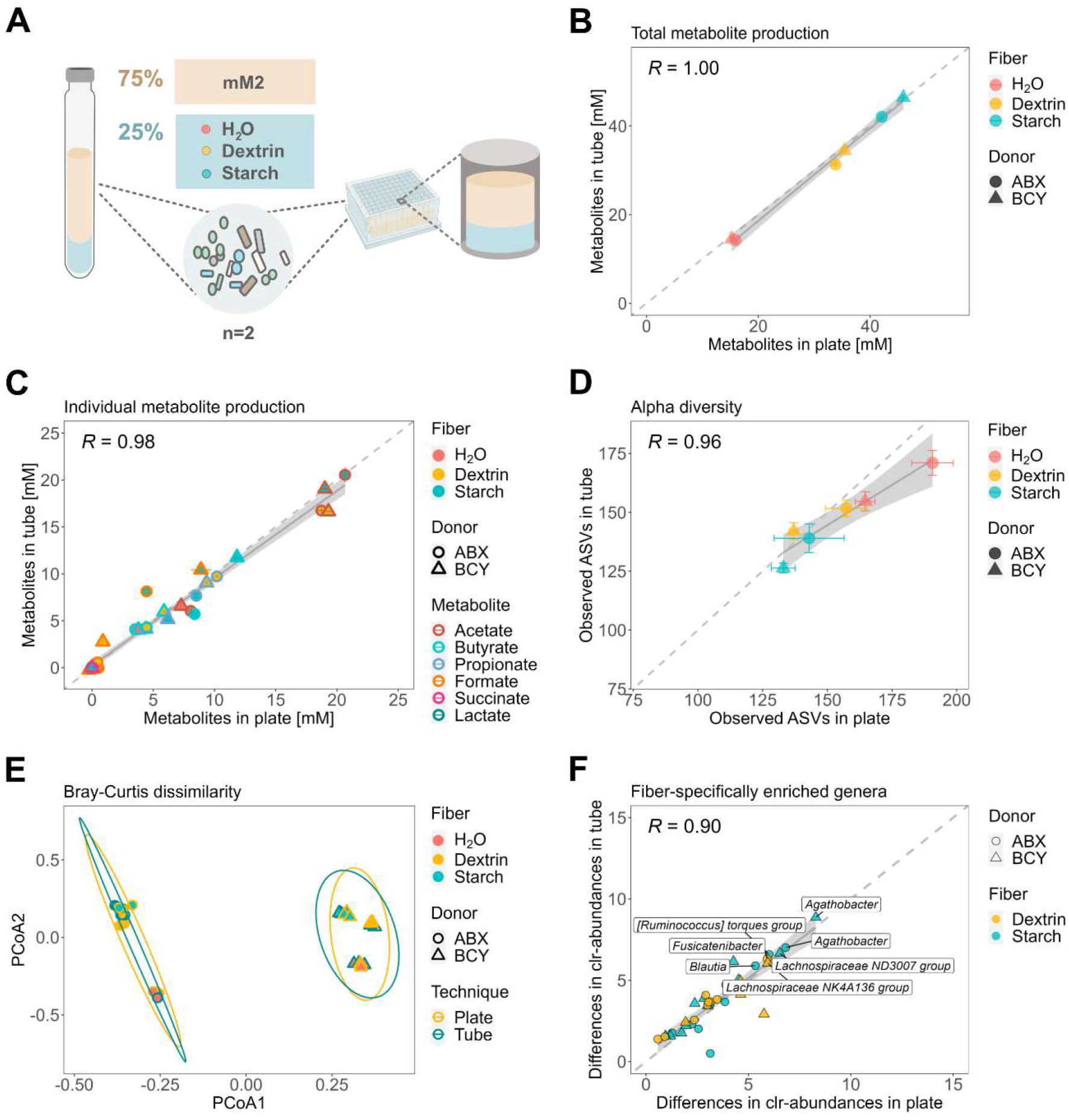
Impact of the cultivation technique (96-deepwell plates vs. gas-tight tubes) on the metabolism and composition of fecal cultures (donor ABX and BCY) cultivated in mM2 for 48 h, 37°C, in the presence of resistant dextrin (3 g/L), soluble starch (3 g/L) or H_2_O as control (technical triplicates). **A)** Experimental setup for comparing the plate- and tube-based techniques. All analyses were performed after 48 h cultivation. **B)** Correlation of the total metabolite production. Total metabolites were calculated by summing the medium-corrected concentrations of organic acids (SCFA and intermediates). **C)** Correlation of individual metabolites. Points represent means of blank-corrected concentrations and bars represent standard deviations. **D)** Correlation of observed ASVs. Points represent means and bars represent standard deviations. **E)** PCoA plot of Bray-Curtis distances with 95% confidence ellipses (calculated separately for each donor) for the two cultivation techniques. **F)** Median clr-differences of taxa (>1% abundance) that are significantly different between each fiber (dextrin, starch) and H_2_O (p ≤0.05). Labels indicate highly enriched taxa (clr-differences >5).

**Figure S4:**
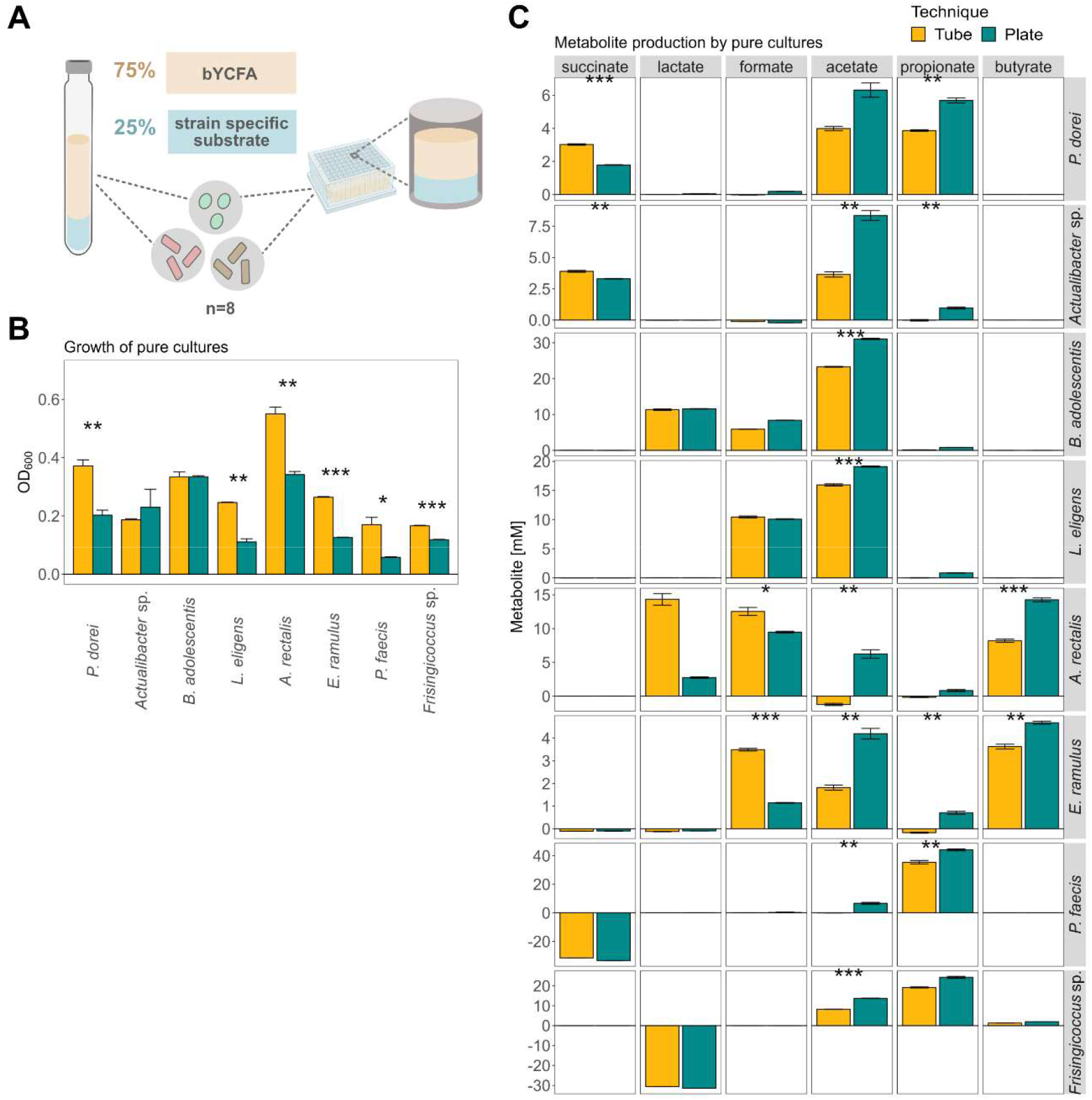
Impact of the cultivation technique (96-deepwell plates vs. gas-tight tubes) on the physiology of pure bacterial strains cultivated in bYCFA for 48 h, 37°C, in the presence of different carbon sources (8 strains; biological triplicates). **A)** Experimental setup for comparing the high-throughput protocol with the Hungate technique using pure cultures. All analyses were performed after 48 h cultivation. **B)** Growth of eight intestinal strains. Bars represent mean endpoint OD_600_ values. **C)** Individual metabolite production. Bars represent mean endpoint concentrations. bYCFA was supplemented with strain-specific substrates (**Table S2**): *E. ramulus* with arabinogalactan, *P. dorei* with pea fiber, *P. faecis* with succinate, *Acutalibacter* sp. with resistant dextrin, *B. adolescentis* with xylan, *A. rectalis* with soluble starch, *Frisingicoccus* sp. with DL-lactate, and *L. eligens* with pectin. Each condition was tested in biological triplicates. Significance was assessed using a t-test with * indicating p≤0.05, ** p≤0.01 and *** p≤ 0.001.

**Figure S5:**
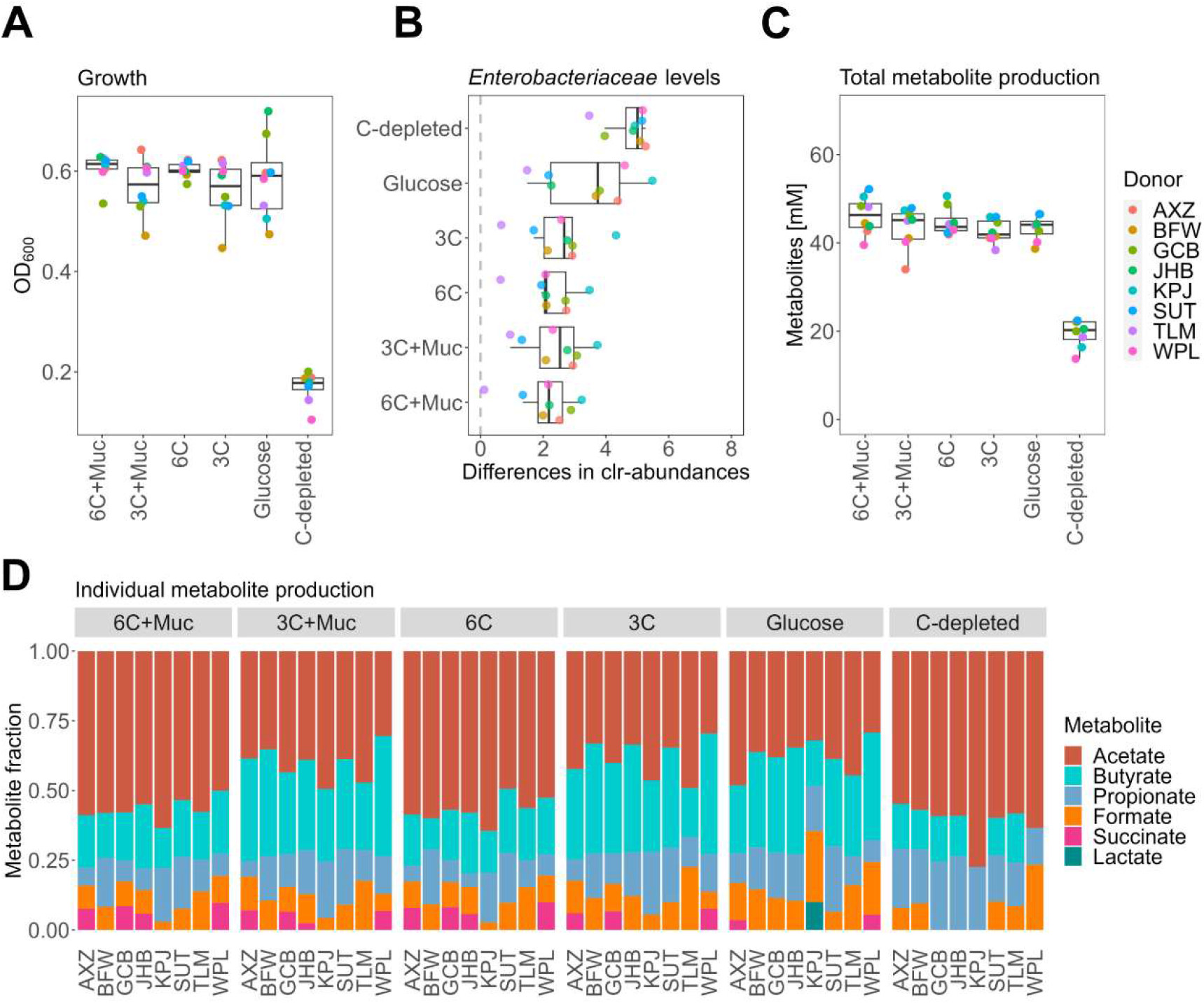
Characteristics of fecal cultures (in bYCFA, 48 h, 37°C, donors n=8) grown in the presence of different C-sources (6C+Muc, 3C+Muc, 6C, 3C and glucose) or in control conditions (C-depleted). All analyses were performed after 48 h cultivation. **A)** Growth (OD_600_) of fecal cultures (200 µL sample; path length ∼5 mm). **B)** Median difference of clr-abundance of *Enterobacteriaceae* in cultures compared to feces. **C)** Total metabolites were calculated by summing the medium-corrected concentrations of organic acids (SCFA and intermediates) in the supernatant of cultures. **C**) Relative metabolite production was calculated by dividing the concentration of each metabolite by the total amount of metabolites produced.

**Figure S6:**
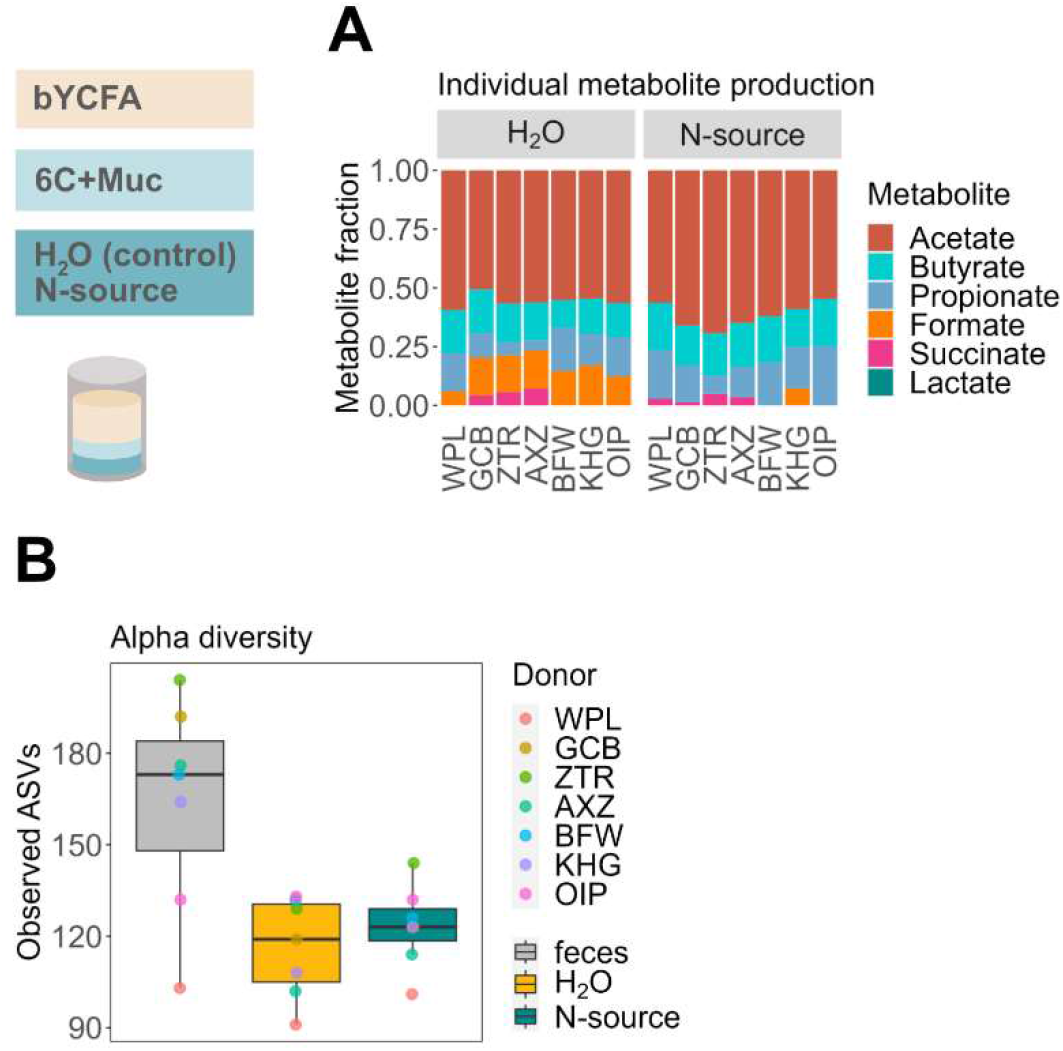
Characteristics of fecal cultures grown in bYCFA with 6C+Muc (48 h, 37°C, donors n=7) supplemented with additional N-source (Amicase and yeast extract) or control (H_2_O) conditions. **A)** Relative metabolite production was calculated by dividing the concentration of each metabolite by the total amount of metabolites produced. **B)** Number of observed ASVs in feces and cultures. All analyses were performed after 48 h on pooled samples from three technical replicates for each donor microbiota.

**Figure S7:**
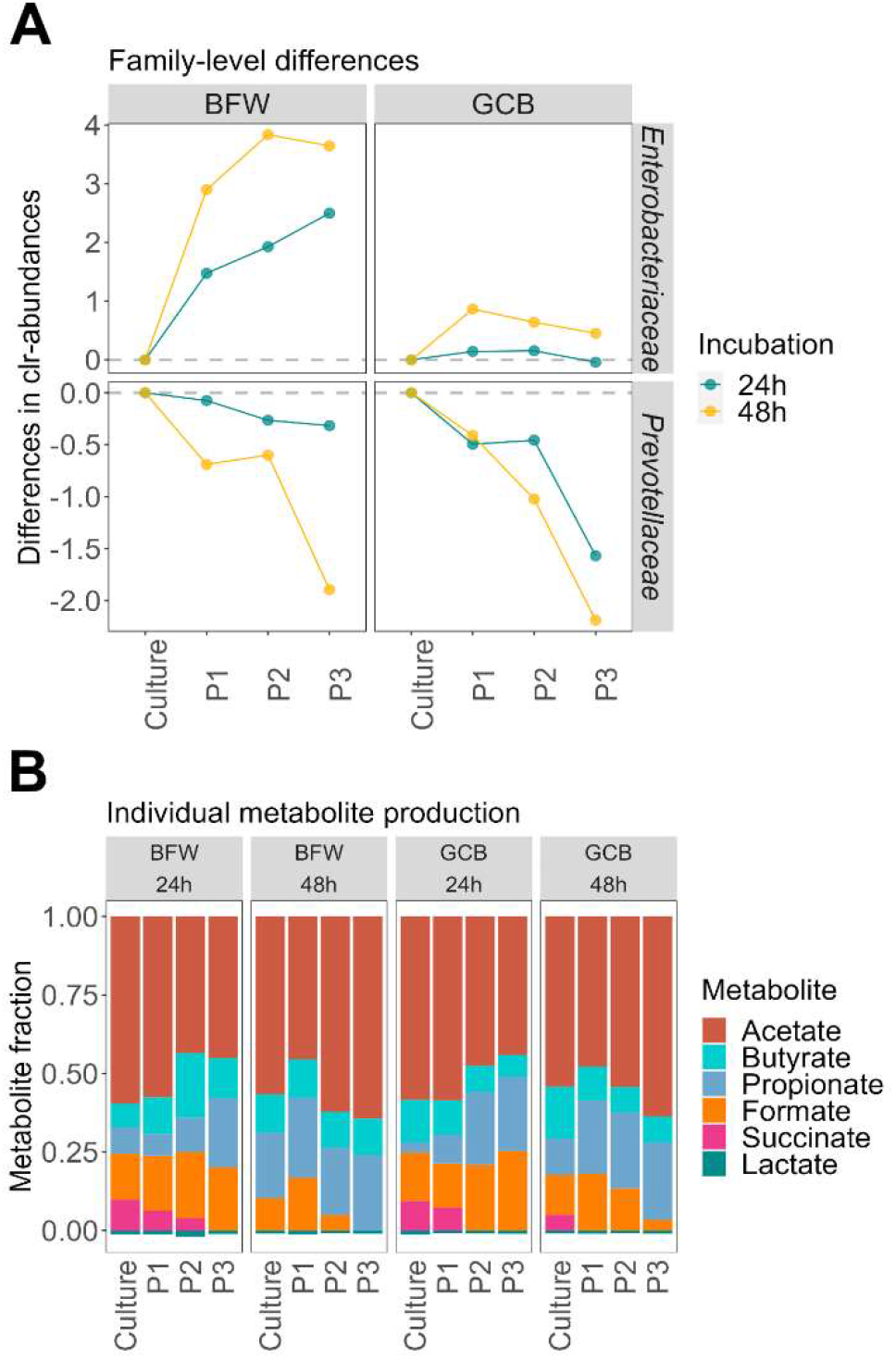
Characteristics of cultures derived from donor microbiota BFW and GCB repeatedly passaged by re-inoculating 1% (3 passages, 24/48 h, 37°C, donors n=2) into bYCFA and 6C+Muc. **A)** Difference of clr-adundance of the families *Enterobacteriaceae* and *Prevotellaceae* between the initial culture and the successive passages (P1, P2, P3). **B)** Relative metabolite production was calculated by dividing the concentration of each metabolite by the total amount of metabolites produced.

**Figure S8:**
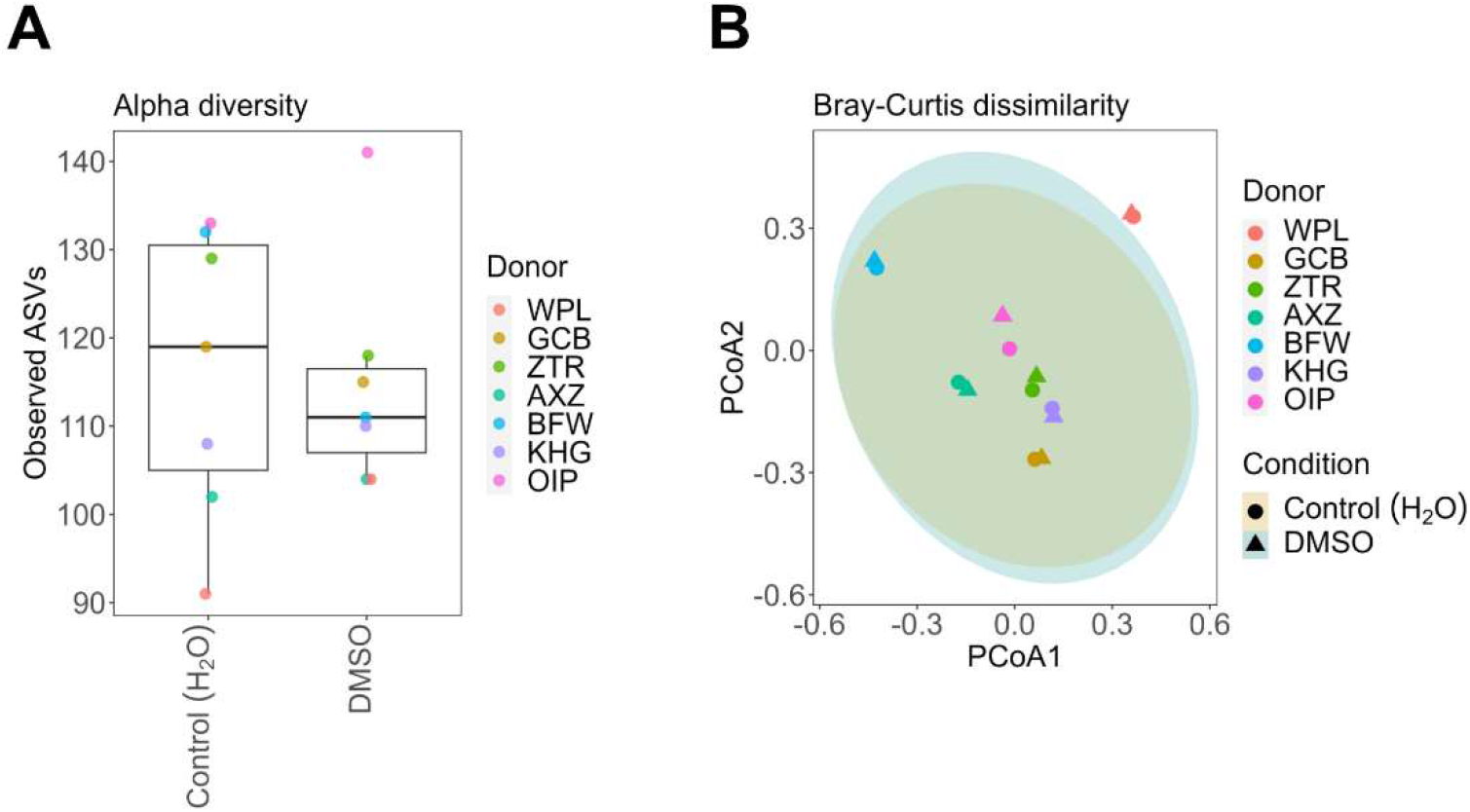
Characteristics of fecal cultures grown in bYCFA with 6C+Muc (48 h, 37°C, donors n=7) and exposed to 0.2 % DMSO compared to control (H_2_O). **A)** Number of observed ASVs in cultures treated with 0.2% DMSO or H_2_O (control). **B)** Bray-Curtis distances between cultures treated with DMSO (0.2%) and H_2_O visualized in a PCoA plot. All analyses were performed after 48 h on pooled samples from three technical replicates for each donor microbiota.

**Figure S9:**
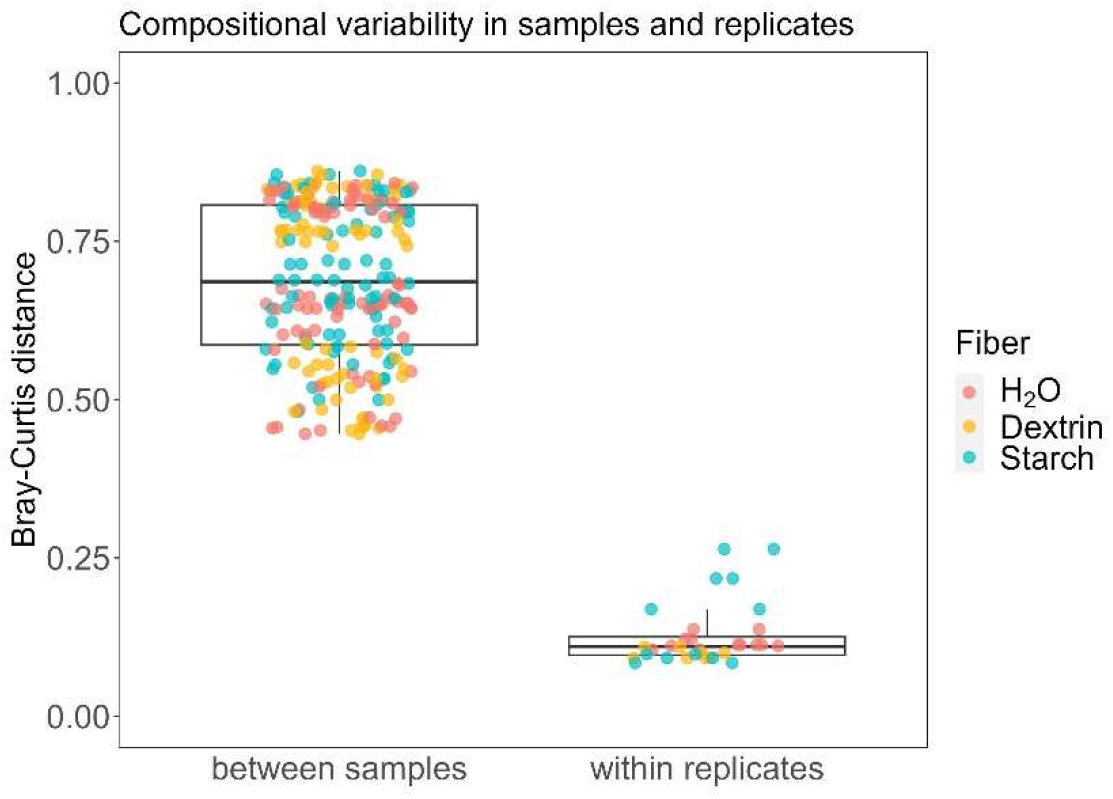
Bray-Curtis distance between samples and within technical replicates of plate-cultured fecal communities (in bYCFA for 48 h, 37°C) in the presence of resistant dextrin (3 g/L), soluble starch (3 g/L) or H_2_O as control (technical triplicates). The displayed samples originate from the experiment comparing the two cultivation techniques (96-deepwell plates vs. gas-tight tubes). Fecal cultures (donor ABX and BCY) were cultivated in bYCFA for 48 h, 37°C, in the presence of resistant dextrin (3 g/L), soluble starch (3 g/L) or H_2_O as control using plates. Distances between samples include distances calculated between cultures under different cultivation conditions or originating from different donor microbiota, while the within replicate distances describe the distance between samples under the same conditions and from the same donor.

## SUPPLEMENTARY TABLES

**Table S1:** Effect of a “loosely” and “strictly” handled chamber on the maximum specific growth rate (µ_max_) and cell density of *F. prausnitzii* DSM 17677 and *B. thetaiotaomicron* DSM 2079. Normality assumption was verified using the Shapiro-Wilk normality test, and significance was determined using unpaired T-test. Significance level was set to *p* < 0.05. The experiments were performed with biological triplicates.

| Strain | Chamber handling | $\mu_{\max}$ (h<sup>-1</sup>) | | | max. cell density (OD<sub>600</sub>) | | |
| --- | --- | --- | --- | --- | --- | --- | --- |
|  |  | Mean | SD | *p* | Mean | SD | *p* |
| *F. prausnitzii*<br>DSM 17677 | H<sub>2</sub>>1.5% | 0.33 | 0.01 | 0.3893 | 0.27 | 0.02 | 0.0023 |
|  | H<sub>2</sub><1.5% | 0.28 | 0.07 |  | 0.06 | 0.04 |  |
| *B. thetaiotaomicron*<br>DSM 2079 | H<sub>2</sub>>1.5% | 0.75 | 0.02 | 0.0145 | 1.03 | 0.01 | 0.0755 |
|  | H<sub>2</sub><1.5% | 0.65 | 0.03 |  | 1.00 | 0.01 |  |

**Table S2:** Bacterial strains used in this study and corresponding carbon source supplemented for growth. Respective fibers were supplemented to reach final 3 g/L, and glucose and organic acids were applied at a concentration of 30 mM.

| Genus | Species | Strain | Culture collection | Supplemented C-source |
| --- | --- | --- | --- | --- |
| *Faecalibacterium* | *prausnitzii* | DSM 17677 | DSMZ | Glucose |
| *Bacteroides* | *thetaiotaomicron* | DSM 2079 | DSMZ | Glucose |
| *Eubacterium* | *ramulus* | PB-SKBSD | PB | Arabinogalactan |
| *Phocaeicola* | *dorei* | PB-SXBFK | PB | Pea fiber |
| *Phascolarctobacterium* | *faecis* | PB-SDVAP | PB | Succinate |
| *Acutalibacter* | sp. | PB-SMJER | PB | Dextrin |
| *Bifidobacterium* | *adolescentis* | PB-SKRVN | PB | Xylan |
| *Agathobacter* | *rectalis* | PB-SWUPP | PB | Soluble starch |
| *Frisingicoccus* | sp. | PB-SKUSI | PB | DL-Lactate |
| *Lachnospira* | *eligans* | PB-SJATG | PB | Pectin |

DSMZ: German Collection of Microorganisms and Cell Culture GmbH; PB: PharmaBiome AG.

**Table S3:** The modular components of the final medium and their composition. The basal medium, heat-stable and heat-sensitive supplements were prepared based on the respective concentration factor and mixed according to % of the final volume.

| Component type | Composition | Concentration factor | % of the final volume |
| --- | --- | --- | --- |
| **Comparison of 96-deep well plates and Hungate tubes using fecal microbiota (Figures 2, S3)** |  |  |  |
| Basal medium | bYCFA, mM2 | 1.33X | 75% |
| Heat-stable supplement | dextrin, starch, H<sub>2</sub>O | 4X | 25% |
| **Comparison of 96-deep well plates and Hungate tubes using pure cultures (Figure S4)** |  |  |  |
| Basal medium | bYCFA | 1.33X | 75% |
| Heat-stable supplement | Arabinogalactan, pea fiber, dextrin, xylan, starch, pectin | 4X | 25% |
| **Optimization of C-source composition to maintain *in vivo*-like communities (Figures 3, S5, S7)** |  |  |  |
| Basal medium | bYCFA | 1.33X | 75% |
| Heat-stable supplement | Glucose, 6C, 6C+Mucin, 3C, 3C+Mucin, H<sub>2</sub>O | 4X | 25% |
| **Optimization of metabolic profile by additional N-source (Figures 3, S6)** |  |  |  |
| Basal medium | bYCFA | 1.33X | 75% |
| Heat-stable supplement | 6C+Mucin | 8X | 12.5% |
| Heat-stable supplement | N-source, H<sub>2</sub>O | 8X | 12.5% |
| **Treatment of fecal cultures with dietary fibers (Figure 4)** |  |  |  |
| Basal medium | bYCFA | 1.33X | 75% |
| Heat-stable supplement | dextrin, starch, H<sub>2</sub>O | 4X | 25% |
| **Treatment of fecal cultures with drugs (Figure 4, S8)** |  |  |  |
| Basal medium | bYCFA | 1.33X | 75% |
| Heat-stable supplement | 6C+Mucin | 8X | 12.5% |
| Heat-sensitive supplement | omeprazole, ciprofloxacin, 5-FU, H<sub>2</sub>O, DMSO | 8X | 12.5% |

## REFERENCES

1. Arumugam, M., Raes, J., Pelletier, E., Paslier, D. Le, Yamada, T., Mende, D.R., Fernandes, G.R., Tap, J., Bruls, T., Batto, J.M., et al. (2011). Enterotypes of the human gut microbiome. Nature 473, 174–180. 10.1038/NATURE09944.

2. Valdes, A.M., Walter, J., Segal, E., and Spector, T.D. (2018). Role of the gut microbiota in nutrition and health. BMJ 361, 36–44. 10.1136/BMJ.K2179.

3. Poeker, S.A., Geirnaert, A., Berchtold, L., Greppi, A., Krych, L., Steinert, R.E., De Wouters, T., and Lacroix, C. (2018). Understanding the prebiotic potential of different dietary fibers using an in vitro continuous adult fermentation model (PolyFermS). Sci. Rep. 8, 1–12. 10.1038/s41598-018-22438-y.

4. Wastyk, H.C., Fragiadakis, G.K., Perelman, D., Dahan, D., Merrill, B.D., Yu, F.B., Topf, M., Gonzalez, C.G., Van Treuren, W., Han, S., et al. (2021). Gut Microbiota-Targeted Diets Modulate Human Immune Status. Cell 184, 4137. 10.1016/J.CELL.2021.06.019.

5. Djekic, D., Shi, L., Brolin, H., Carlsson, F., Särnqvist, C., Savolainen, O., Cao, Y., Bäckhed, F., Tremaroli, V., Landberg, R., et al. (2020). Effects of a Vegetarian Diet on Cardiometabolic Risk Factors, Gut Microbiota, and Plasma Metabolome in Subjects With Ischemic Heart Disease: A Randomized, Crossover Study. J. Am. Hear. Assoc. Cardiovasc. Cerebrovasc. Dis. 9, e016518. 10.1161/JAHA.120.016518.

6. Jackson, M.A., Verdi, S., Maxan, M.E., Shin, C.M., Zierer, J., Bowyer, R.C.E., Martin, T., Williams, F.M.K., Menni, C., Bell, J.T., et al. (2018). Gut microbiota associations with common diseases and prescription medications in a population-based cohort. Nat. Commun. 9, 2655. 10.1038/s41467-018-05184-7.

7. Palleja, A., Mikkelsen, K.H., Forslund, S.K., Kashani, A., Allin, K.H., Nielsen, T., Hansen, T.H., Liang, S., Feng, Q., Zhang, C., et al. (2018). Recovery of gut microbiota of healthy adults following antibiotic exposure. Nat. Microbiol. 3, 1255–1265. 10.1038/s41564-018-0257-9.

8. Klünemann, M., Andrejev, S., Blasche, S., Mateus, A., Phapale, P., Devendran, S., Vappiani, J., Simon, B., Scott, T.A., Kafkia, E., et al. (2021). Bioaccumulation of therapeutic drugs by human gut bacteria. Nature 597, 533–538. 10.1038/s41586-021-03891-8.

9. Hashimoto, T., Perlot, T., Rehman, A., Trichereau, J., Ishiguro, H., Paolino, M., Sigl, V., Hanada, T., Hanada, R., Lipinski, S., et al. (2012). ACE2 links amino acid malnutrition to microbial ecology and intestinal inflammation. Nature 487, 477–481. 10.1038/nature11228.

10. Tropini, C., Moss, E.L., Merrill, B.D., Ng, K.M., Higginbottom, S.K., Casavant, E.P., Gonzalez, C.G., Fremin, B., Bouley, D.M., Elias, J.E., et al. (2018). Transient Osmotic Perturbation Causes Long-Term Alteration to the Gut Microbiota. Cell 173, 1742–1754. 10.1016/J.CELL.2018.05.008.

11. Roager, H.M., Hansen, L.B.S., Bahl, M.I., Frandsen, H.L., Carvalho, V., Gøbel, R.J., Dalgaard, M.D., Plichta, D.R., Sparholt, M.H., Vestergaard, H., et al. (2016). Colonic transit time is related to bacterial metabolism and mucosal turnover in the gut. Nat. Microbiol. 1, 1–9. 10.1038/nmicrobiol.2016.93.

12. Murga-Garrido, S.M., Hong, Q., Cross, T.W.L., Hutchison, E.R., Han, J., Thomas, S.P., Vivas, E.I., Denu, J., Ceschin, D.G., Tang, Z.Z., et al. (2021). Gut microbiome variation modulates the effects of dietary fiber on host metabolism. Microbiome 9, 117. 10.1186/S40168-021-01061-6.

13. Hungate, R.E. (1969). Chapter IV A Roll Tube Method for Cultivation of Strict Anaerobes. Methods Microbiol. 3, 117–132. 10.1016/S0580-9517(08)70503-8.

14. Hungate, R.E. (1944). Studies on Cellulose Fermentation: I. The Culture and Physiology of an Anaerobic Cellulose-digesting Bacterium. J. Bacteriol. 48, 499–513. 10.1128/JB.48.5.499-513.1944.

15. Kundra, P., Geirnaert, A., Pugin, B., Morales Martinez, P., Lacroix, C., and Greppi, A. (2022). Healthy adult gut microbiota sustains its own vitamin B12 requirement in an in vitro batch fermentation model. Front. Nutr. 9, 2940. 10.3389/FNUT.2022.1070155.

16. Sandberg, J., Kovatcheva-Datchary, P., Björck, I., Bäckhed, F., and Nilsson, A. (2019). Abundance of gut Prevotella at baseline and metabolic response to barley prebiotics. Eur. J. Nutr. 58, 2365–2376. 10.1007/S00394-018-1788-9.

17. Walker, A.W., Ince, J., Duncan, S.H., Webster, L.M., Holtrop, G., Ze, X., Brown, D., Stares, M.D., Scott, P., Bergerat, A., et al. (2011). Dominant and diet-responsive groups of bacteria within the human colonic microbiota. ISME J. 5, 220–230. 10.1038/ismej.2010.118.

18. Cantu-Jungles, T.M., Bulut, N., Chambry, E., Ruthes, A., Iacomini, M., Keshavarzian, A., Johnson, T.A., and Hamaker, B.R. (2021). Dietary fiber hierarchical specificity: The missing link for predictable and strong shifts in gut bacterial communities. MBio 12, e01028–21. 10.1128/MBIO.01028-21.

19. Hjorth, M.F., Christensen, L., Larsen, T.M., Roager, H.M., Krych, L., Kot, W., Nielsen, D.S., Ritz, C., and Astrup, A. (2020). Pretreatment Prevotella-to-Bacteroides ratio and salivary amylase gene copy number as prognostic markers for dietary weight loss. Am. J. Clin. Nutr. 111, 1079–1086. doi.org/10.1093/ajcn/nqaa007.

20. Hall, A.B., Yassour, M., Sauk, J., Garner, A., Jiang, X., Arthur, T., Lagoudas, G.K., Vatanen, T., Fornelos, N., Wilson, R., et al. (2017). A novel Ruminococcus gnavus clade enriched in inflammatory bowel disease patients. Genome Med. 9, 1–12. 10.1186/S13073-017-0490-5.

21. Karcher, N., Pasolli, E., Asnicar, F., Huang, K.D., Tett, A., Manara, S., Armanini, F., Bain, D., Duncan, S.H., Louis, P., et al. (2020). Analysis of 1321 Eubacterium rectale genomes from metagenomes uncovers complex phylogeographic population structure and subspecies functional adaptations. Genome Biol. 21, 138. 10.1186/S13059-020-02042-Y.

22. Sorbara, M.T., Littmann, E.R., Fontana, E., Moody, T.U., Kohout, C.E., Gjonbalaj, M., Eaton, V., Seok, R., Leiner, I.M., and Pamer, E.G. (2020). Functional and Genomic Variation between Human-Derived Isolates of Lachnospiraceae Reveals Inter- and Intra-Species Diversity. Cell Host Microbe 28, 134–146. 10.1016/J.CHOM.2020.05.005.

23. Otaru, N., Ye, K., Mujezinovic, D., Berchtold, L., Constancias, F., Cornejo, F.A., Krzystek, A., de Wouters, T., Braegger, C., Lacroix, C., et al. (2021). GABA Production by Human Intestinal Bacteroides spp.: Prevalence, Regulation, and Role in Acid Stress Tolerance. Front. Microbiol. 12, 656895. 10.3389/FMICB.2021.656895.

24. Anthamatten, L., Bieberstein, P.R. von, Thabuis, C., Menzi, C., Reichlin, M., Meola, M., Rodriguez, B., Cordero, O.X., Lacroix, C., Wouters, T. de, et al. (2023). Mapping gut bacteria into functional niches reveals the ecological structure of human gut microbiomes. bioRxiv, 2023.07.04.547750. 10.1101/2023.07.04.547750.

25. Aranda-Díaz, A., Ng, K.M., Thomsen, T., Real-Ramírez, I., Dahan, D., Dittmar, S., Gonzalez, C.G., Chavez, T., Vasquez, K.S., Nguyen, T.H., et al. (2022). Establishment and characterization of stable, diverse, fecal-derived in vitro microbial communities that model the intestinal microbiota. Cell Host Microbe 30, 260–272. 10.1016/J.CHOM.2021.12.008.

26. Li, L., Abou-Samra, E., Ning, Z., Zhang, X., Mayne, J., Wang, J., Cheng, K., Walker, K., Stintzi, A., and Figeys, D. (2019). An in vitro model maintaining taxon-specific functional activities of the gut microbiome. Nat. Commun. 10, 4146. 10.1038/s41467-019-12087-8.

27. Javdan, B., Lopez, J.G., Chankhamjon, P., Lee, Y.C.J., Hull, R., Wu, Q., Wang, X., Chatterjee, S., and Donia, M.S. (2020). Personalized mapping of drug metabolism by the human gut microbiome. Cell 181, 1661. 10.1016/J.CELL.2020.05.001.

28. Li, L., Ning, Z., Zhang, X., Mayne, J., Cheng, K., Stintzi, A., and Figeys, D. (2020). RapidAIM: A culture- And metaproteomics-based Rapid Assay of Individual Microbiome responses to drugs. Microbiome 8, 1–16. 10.1186/S40168-020-00806-Z.

29. Douglas S. Auld, P.D., Peter A. Coassin, B.S., Nathan P. Coussens, P.D., Hensley, P., Klumpp-Thomas, C., Michael, S., G. Sitta Sittampalam, P.D., O. Joseph Trask, B.S., Bridget K. Wagner, P.D., Jeffrey R. Weidner, P.D., et al. (2020). Microplate Selection and Recommended Practices in High-throughput Screening and Quantitative Biology (Eli Lilly & Company and the National Center for Advancing Translational Sciences).

30. Miyazaki, K., Martin, J.C., Marinsek-Logar, R., and Flint, H.J. (1997). Degradation and Utilization of Xylans by the Rumen AnaerobePrevotella bryantii(formerlyP. ruminicolasubsp.brevis) B14. Anaerobe 3, 373–381. 10.1006/ANAE.1997.0125.

31. Duncan, S.H., Hold, G.L., Harmsen, H.J.M., Stewart, C.S., and Flint, H.J. (2002). Growth requirements and fermentation products of Fusobacterium prausnitzii, and a proposal to reclassify it as Faecalibacterium prausnitzii gen. nov., comb. nov. Int. J. Syst. Evol. Microbiol. 52, 2141–2146. 10.1099/00207713-52-6-2141.

32. Kohn, R.A., and Dunlap, T.F. (1998). Calculation of the buffering capacity of bicarbonate in the rumen and in vitro. J. Anim. Sci. 76, 1702–1709. 10.2527/1998.7661702X.

33. Macfarlane, G.T., Macfarlane, S., and Gibson, G.R. (1998). Validation of a three-stage compound continuous culture system for investigating the effect of retention time on the ecology and metabolism of bacteria in the human colon. Microb. Ecol. 35, 180–187. 10.1007/S002489900072.

34. Isenring, J., Bircher, L., Geirnaert, A., and Lacroix, C. (2023). In vitro human gut microbiota fermentation models: opportunities, challenges, and pitfalls. Microbiome Res. Reports 2, 2. 10.20517/MRR.2022.15.

35. Walker, A.W., Duncan, S.H., Carol McWilliam Leitch, E., Child, M.W., and Flint, H.J. (2005). pH and peptide supply can radically alter bacterial populations and short-chain fatty acid ratios within microbial communities from the human colon. Appl. Environ. Microbiol. 71, 3692–3700. 10.1128/AEM.71.7.3692-3700.2005.

36. Cummings, J.H., Pomare, E.W., Branch, H.W.J., Naylor, E., and Macfarlane, G.T. (1987). Short chain fatty acids in human large intestine, portal, hepatic and venous blood. Gut 28, 1221–1227. 10.1136/gut.28.10.1221.

37. Guerin-Deremaux, L., Ringard, F., Desailly, F., and Wils, D. (2010). Effects of a soluble dietary fibre NUTRIOSE® on colonic fermentation and excretion rates in rats. Nutr. Res. Pract. 4, 470–476. 10.4162/NRP.2010.4.6.470.

38. Louis, P., Scott, K.P., Duncan, S.H., and Flint, H.J. (2007). Understanding the effects of diet on bacterial metabolism in the large intestine. J. Appl. Microbiol. 102, 1197–1208. 10.1111/J.1365-2672.2007.03322.X.

39. Venkataraman, A., Sieber, J.R., Schmidt, A.W., Waldron, C., Theis, K.R., and Schmidt, T.M. (2016). Variable responses of human microbiomes to dietary supplementation with resistant starch. Microbiome 4, 1–9. 10.1186/S40168-016-0178-X.

40. Baxter, N.T., Schmidt, A.W., Venkataraman, A., Kim, K.S., Waldron, C., and Schmidt, T.M. (2019). Dynamics of human gut microbiota and short-chain fatty acids in response to dietary interventions with three fermentable fibers. MBio 10, e02566–18. 10.1128/MBIO.02566-18.

41. Barber, C., Sabater, C., Ávila-Gálvez, M.Á., Vallejo, F., Bendezu, R.A., Guérin-Deremaux, L., Guarner, F., Espín, J.C., Margolles, A., and Azpiroz, F. (2022). Effect of Resistant Dextrin on Intestinal Gas Homeostasis and Microbiota. Nutrients 14, 4611. 10.3390/NU14214611.

42. Vital, M., Howe, A., Bergeron, N., Krauss, R.M., Jansson, J.K., and Tiedje, J.M. (2018). Metagenomic Insights into the Degradation of Resistant Starch by Human Gut Microbiota. Appl. Environ. Microbiol. 84, e.01562–18. 10.1128/AEM.01562-18.

43. Vich Vila, A., Collij, V., Sanna, S., Sinha, T., Imhann, F., Bourgonje, A.R., Mujagic, Z., Jonkers, D.M.A.E., Masclee, A.A.M., Fu, J., et al. (2020). Impact of commonly used drugs on the composition and metabolic function of the gut microbiota. Nat. Commun. 11, 362. 10.1038/s41467-019-14177-z.

44. van Kessel, S.P., Frye, A.K., El-Gendy, A.O., Castejon, M., Keshavarzian, A., van Dijk, G., and El Aidy, S. (2019). Gut bacterial tyrosine decarboxylases restrict levels of levodopa in the treatment of Parkinson’s disease. Nat. Commun. 10, 310. 10.1038/s41467-019-08294-y.

45. Yuan, L., Zhang, S., Li, H., Yang, F., Mushtaq, N., Ullah, S., Shi, Y., An, C., and Xu, J. (2018). The influence of gut microbiota dysbiosis to the efficacy of 5-Fluorouracil treatment on colorectal cancer. Biomed. Pharmacother. 108, 184–193. 10.1016/J.BIOPHA.2018.08.165.

46. Zimmermann, M., Zimmermann-Kogadeeva, M., Wegmann, R., and Goodman, A.L. (2019). Mapping human microbiome drug metabolism by gut bacteria and their genes. Nature 570, 462–467. 10.1038/s41586-019-1291-3.

47. Maier, L., Pruteanu, M., Kuhn, M., Zeller, G., Telzerow, A., Anderson, E.E., Brochado, A.R., Fernandez, K.C., Dose, H., Mori, H., et al. (2018). Extensive impact of non-antibiotic drugs on human gut bacteria. Nature 555, 623–628. 10.1038/nature25979.

48. LaCourse, K.D., Zepeda-Rivera, M., Kempchinsky, A.G., Baryiames, A., Minot, S.S., Johnston, C.D., and Bullman, S. (2022). The cancer chemotherapeutic 5-fluorouracil is a potent Fusobacterium nucleatum inhibitor and its activity is modified by intratumoral microbiota. Cell Rep. 41. 10.1016/J.CELREP.2022.111625.

49. Stewardson, A.J., Gaïa, N., François, P., Malhotra-Kumar, S., Delémont, C., Martinez de Tejada, B., Schrenzel, J., Harbarth, S., Lazarevic, V., Vervoort, J., et al. (2015). Collateral damage from oral ciprofloxacin versus nitrofurantoin in outpatients with urinary tract infections: a culture-free analysis of gut microbiota. Clin. Microbiol. Infect. 21, 344.e1. 10.1016/J.CMI.2014.11.016.

50. Li, H.L., Lu, L., Wang, X.S., Qin, L.Y., Wang, P., Qiu, S.P., Wu, H., Huang, F., Zhang, B.B., Shi, H.L., et al. (2017). Alteration of gut microbiota and inflammatory cytokine/chemokine profiles in 5-fluorouracil induced intestinal mucositis. Front. Cell. Infect. Microbiol. 7, 455. 10.3389/FCIMB.2017.00455.

51. Hamouda, N., Sano, T., Oikawa, Y., Ozaki, T., Shimakawa, M., Matsumoto, K., Amagase, K., Higuchi, K., and Kato, S. (2017). Apoptosis, Dysbiosis and Expression of Inflammatory Cytokines are Sequential Events in the Development of 5-Fluorouracil-Induced Intestinal Mucositis in Mice. Basic Clin. Pharmacol. Toxicol. 121, 159–168. 10.1111/BCPT.12793.

52. Carvalho, R., Vaz, A., Pereira, F.L., Dorella, F., Aguiar, E., Chatel, J.M., Bermudez, L., Langella, P., Fernandes, G., Figueiredo, H., et al. (2018). Gut microbiome modulation during treatment of mucositis with the dairy bacterium Lactococcus lactis and recombinant strain secreting human antimicrobial PAP. Sci. Rep. 8, 15072. 10.1038/s41598-018-33469-w.

53. Jackson, M.A., Goodrich, J.K., Maxan, M.E., Freedberg, D.E., Abrams, J.A., Poole, A.C., Sutter, J.L., Welter, D., Ley, R.E., Bell, J.T., et al. (2016). Proton pump inhibitors alter the composition of the gut microbiota. Gut 65, 749–756. 10.1136/GUTJNL-2015-310861.

54. Clavel, T., Horz, H.P., Segata, N., and Vehreschild, M. (2022). Next steps after 15 stimulating years of human gut microbiome research. Microb. Biotechnol. 15, 164–175. 10.1111/1751-7915.13970.

55. Hitch, T.C.A., Afrizal, A., Riedel, T., Kioukis, A., Haller, D., Lagkouvardos, I., Overmann, J., and Clavel, T. (2021). Recent advances in culture-based gut microbiome research. Int. J. Med. Microbiol. 311, 151485. 10.1016/J.IJMM.2021.151485.

56. Goodman, A.L., Kallstrom, G., Faith, J.J., Reyes, A., Moore, A., Dantas, G., and Gordon, J.I. (2011). Extensive personal human gut microbiota culture collections characterized and manipulated in gnotobiotic mice. Proc. Natl. Acad. Sci. U. S. A. 108, 6252–6257. 10.1073/PNAS.1102938108.

57. Berg, M., Undisz, K., Thiericke, R., Zimmermann, P., Moore, T., and Posten, C. (2001). Evaluation of Liquid Handling Conditions in Microplates. SLAS Discov. 6, 47–56. 10.1177/108705710100600107.

58. Fallingborg, J. (1999). Intraluminal pH of the human gastrointestinal tract. Dan. Med. Bull. 46, 183–196.

59. Koziolek, M., Grimm, M., Becker, D., Iordanov, V., Zou, H., Shimizu, J., Wanke, C., Garbacz, G., and Weitschies, W. (2015). Investigation of pH and Temperature Profiles in the GI Tract of Fasted Human Subjects Using the Intellicap(®) System. J. Pharm. Sci. 104, 2855–2863. 10.1002/JPS.24274.

60. Fischbach, M.A., and Sonnenburg, J.L. (2011). Eating For Two: How Metabolism Establishes Interspecies Interactions in the Gut. Cell Host Microbe 10, 336. 10.1016/J.CHOM.2011.10.002.

61. Macy, J.M., Ljungdahl, L.G., and Gottschalk, G. (1978). Pathway of Succinate and Propionate Formation in Bacteroides fragilis. J. Bacteriol. 134, 84–91. 10.1128/JB.134.1.84-91.1978.

62. Smith, N.W., Shorten, P.R., Altermann, E.H., Roy, N.C., and McNabb, W.C. (2019). Hydrogen cross-feeders of the human gastrointestinal tract. Gut Microbes 10, 270–288. 10.1080/19490976.2018.1546522.

63. Van Lingen, H.J., Plugge, C.M., Fadel, J.G., Kebreab, E., Bannink, A., and Dijkstra, J. (2016). Thermodynamic Driving Force of Hydrogen on Rumen Microbial Metabolism: A Theoretical Investigation. PLoS One 11. 10.1371/JOURNAL.PONE.0161362.

64. Modesto, A., Cameron, N.R., Varghese, C., Peters, N., Stokes, B., Phillips, A., Bissett, I., and O’Grady, G. (2022). Meta-Analysis of the Composition of Human Intestinal Gases. Dig. Dis. Sci. 67, 3842–3859. 10.1007/S10620-021-07254-1.

65. Chauveau, A., Treyer, A., Geirnaert, A., Bircher, L., Babst, A., Abegg, V.F., Simões-Wüst, A.P., Lacroix, C., Potterat, O., and Hamburger, M. (2023). Intestinal permeability and gut microbiota interactions of pharmacologically active compounds in valerian and St. John’s wort. Biomed. Pharmacother. 162, 114652. 10.1016/J.BIOPHA.2023.114652.

66. Shetty, S.A., Kuipers, B., Atashgahi, S., Aalvink, S., Smidt, H., and de Vos, W.M. (2022). Inter-species Metabolic Interactions in an In-vitro Minimal Human Gut Microbiome of Core Bacteria. npj Biofilms Microbiomes 8, 21. 10.1038/s41522-022-00275-2.

67. Liu, Y., Gibson, G.R., and Walton, G.E. (2016). An In Vitro Approach to Study Effects of Prebiotics and Probiotics on the Faecal Microbiota and Selected Immune Parameters Relevant to the Elderly. PLoS One 11, e0162604. 10.1371/JOURNAL.PONE.0162604.

68. Gibbons, S.M., Gurry, T., Lampe, J.W., Chakrabarti, A., Dam, V., Everard, A., Goas, A., Gross, G., Kleerebezem, M., Lane, J., et al. (2022). Perspective: Leveraging the Gut Microbiota to Predict Personalized Responses to Dietary, Prebiotic, and Probiotic Interventions. Adv. Nutr. 13, 1450–1461. 10.1093/ADVANCES/NMAC075.

69. Relizani, K., Le Corf, K., Kropp, C., Martin-Rosique, R., Kissi, D., Déjean, G., Bruno, L., Martinez, C., Rawadi, G., Elustondo, F., et al. (2022). Selection of a novel strain of Christensenella minuta as a future biotherapy for Crohn’s disease. Sci. Rep. 12. 10.1038/S41598-022-10015-3.

70. Bircher, L., Geirnaert, A., Hammes, F., Lacroix, C., and Schwab, C. (2018). Effect of cryopreservation and lyophilization on viability and growth of strict anaerobic human gut microbes. Microb. Biotechnol. 11, 721–733. 10.1111/1751-7915.13265.

71. Caporaso, J.G., Lauber, C.L., Walters, W.A., Berg-Lyons, D., Huntley, J., Fierer, N., Owens, S.M., Betley, J., Fraser, L., Bauer, M., et al. (2012). Ultra-high-throughput microbial community analysis on the Illumina HiSeq and MiSeq platforms. ISME J. 6, 1621–1624. 10.1038/ismej.2012.8.

72. Caporaso, J.G., Lauber, C.L., Walters, W.A., Berg-Lyons, D., Lozupone, C.A., Turnbaugh, P.J., Fierer, N., and Knight, R. (2011). Global patterns of 16S rRNA diversity at a depth of millions of sequences per sample. Proc. Natl. Acad. Sci. U. S. A. 108, 4516–4522. 10.1073/PNAS.1000080107.

73. Walters, W., Hyde, E.R., Berg-Lyons, D., Ackermann, G., Humphrey, G., Parada, A., Gilbert, J.A., Jansson, J.K., Caporaso, J.G., Fuhrman, J.A., et al. (2015). Improved Bacterial 16S rRNA Gene (V4 and V4-5) and Fungal Internal Transcribed Spacer Marker Gene Primers for Microbial Community Surveys. mSystems 1. 10.1128/MSYSTEMS.00009-15.

74. Constancias, F., and Mahé, F. (2022). fconstancias/metabaRpipe: v0.9 (v0.9). Zenodo. 10.5281/zenodo.6423397.

75. Callahan, B.J., McMurdie, P.J., Rosen, M.J., Han, A.W., Johnson, A.J.A., and Holmes, S.P. (2016). DADA2: High-resolution sample inference from Illumina amplicon data. Nat. Methods 13, 581–583. 10.1038/nmeth.3869.

76. Quast, C., Pruesse, E., Yilmaz, P., Gerken, J., Schweer, T., Yarza, P., Peplies, J., and Glöckner, F.O. (2013). The SILVA ribosomal RNA gene database project: improved data processing and web-based tools. Nucleic Acids Res. 41, D590–D596. 10.1093/NAR/GKS1219.

77. Yilmaz, P., Parfrey, L.W., Yarza, P., Gerken, J., Pruesse, E., Quast, C., Schweer, T., Peplies, J., Ludwig, W., and Glöckner, F.O. (2014). The SILVA and “All-species Living Tree Project (LTP)” taxonomic frameworks. Nucleic Acids Res. 42, D643–D648. 10.1093/NAR/GKT1209.

78. McMurdie, P.J., and Holmes, S. (2013). phyloseq: An R Package for Reproducible Interactive Analysis and Graphics of Microbiome Census Data. PLoS One 8, e61217. 10.1371/JOURNAL.PONE.0061217.

79. Fernandes, A.D., Reid, J.N.S., Macklaim, J.M., McMurrough, T.A., Edgell, D.R., and Gloor, G.B. (2014). Unifying the analysis of high-throughput sequencing datasets: Characterizing RNA-seq, 16S rRNA gene sequencing and selective growth experiments by compositional data analysis. Microbiome 2, 1–13. 10.1186/2049-2618-2-15.

