## Supplementary file 1 for "A flexible high-throughput cultivation protocol to assess the response of individuals’ gut microbiota to diet-, drug-, and host-related factors"

### Supplementary file 1: Protocol for the high-throughput cultivation of gut microbiota and microbes in an anaerobic chamber

#### *A) Purpose and Principle*

This protocol was developed to test the responses of individuals' gut microbiota and pure cultures to various factors of the intestine related to diet (e.g., fibers, proteins, vitamins), host physiology (*e.g.*, pH, oxidative stress, bile salts) or drugs (*e.g.*, antibiotic and non-antibiotic medications).

The protocol consists of several steps, starting with preparation of the medium and solutions (**Step 1**) several days before the experiment. This is followed by setting up the anaerobic chamber (**Step 2**) and allowing the medium to equilibrate (**Step 3**) on the day before the experiment. The culture conditions are then finalized (**Step 4 to 6**), and the microbes are cultivated (**Step 7**) on the day of the experiment. Finally, sample preparation and analysis are described (**Step 8**). On the day of the experiment (**Step 4 to Step 7**), it is possible to work alone. However, if the number of fecal donors and/or conditions to be tested is high, it should ideally be done by two people in parallel to ensure that inoculation is performed immediately after the pH adjustment of the basal medium.

The medium comprises modular parts (the basal medium and heat-stable/sensitive supplements) that are separately customizable, providing high flexibility. Here, the step-by-step protocol was written based on the example of testing fecal cultures' responses to drugs. In this specific example, the medium is composed of 75% bYCFA (basal medium, 1.33X concentrated), 12.5% 6C+Muc (heat-stable supplement, 8X concentrated), and 12.5% drug solutions (heat-sensitive supplement, 8X concentrated). Depending on the research question, the experimental design can be simplified or expanded to other medium compositions, treatments (*e.g.,* dietary fibers), and physicochemical conditions (*e.g.,* pH).

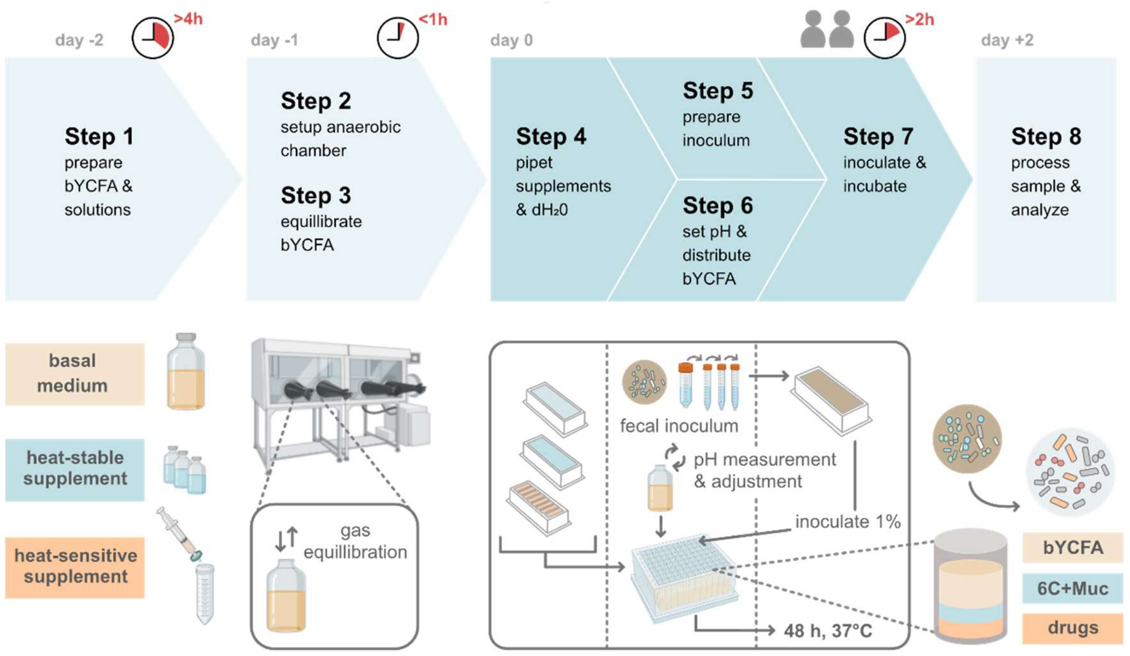


**Figure 1:** Overview of the different Steps of the high-throughput protocol for the cultivation of gut microbiota and microbes.

##### *B) Reagent composition and preparation*

*Note: Aerobic solutions can be prepared directly on the bench. The anaerobic medium and anaerobic heat-stable solutions are boiled and when cooled down, either $CO_2$ or $N_2$ can be used (interchangeably, depending on availability) to help remove $O_2$ traces. However, consider that flushing with $CO_2$ (preferred for bicarbonate-buffered media) acidifies the solution, while $N_2$ does not (preferred for non-buffered solutions). After transfer to the anaerobic chamber, the medium pH will change again depending on the $CO_2$ concentration in the chamber.*

**#1 Vitamin solution (aerobic):** Dissolve 10 mg biotin, 10 mg cyanocobalamin, 30 mg *p*-aminobenzoic acid, 50 mg folic acid, and 150 mg pyridoxamine in 100 mL deionized water ($dH_2O$). Cover aliquots with aluminum foil and store them at 4°C (max. 3 months) or -20°C (long-term storage).

**#2 Mineral solution I (aerobic):** Dissolve 3.5 g $K_2HPO_4$ in 1 L $dH_2O$. Store at 4°C.

**#3 Mineral solution II (aerobic):** Dissolve 3.5 g $KH_2PO_4$, 6 g NaCl, 6 g $(NH_4)_2SO_4$, 0.6 g $MgSO_4$, and 0.6 g $CaCl_2$ in 1 L $dH_2O$. Store at 4°C.

**#4 Hemin stock solution (aerobic):** Dissolve 1.25 g hemin from bovine in 5 mL NaOH 5 M and add 20 mL $dH_2O$. Cover with aluminum foil and store at 4°C (max. 3 months).

**#5 Volatile fatty acid solution (aerobic):** In a chemical hood, on ice, mix 17 mL acetic acid (>99.5%), 6 mL propionic acid (>99.5%), 1 mL isovaleric acid (>99%), 1 mL isobutyric acid (>99%), and 1 mL valeric acid (>99%), and complement with 24 mL of 5 M NaOH. Store at room temperature.

**#6 Resazurin stock solution (aerobic):** Dissolve 100 mg resazurin sodium salt in 100 mL $dH_2O$. Store at 4°C.

**#7 bYCFA basal media (1.33X concentrated, anaerobic):** The cultivation medium is prepared with a concentration factor (here 1.33X) to account for further addition of (heat-stable/sensitive) supplements (here C-source mix and drug solutions). In an Erlenmeyer flask, mix all nitrogen sources (1 g amicase, 1.25 g yeast extract, 0.5 g meat extract) with 0.1 mL vitamin solution (**Reagent #1**), 150 mL mineral solution I (**Reagent #2**), 150 mL mineral solution II (**Reagent #3**), 0.2 mL hemin stock solution (**Reagent #4**) and 6.2 mL volatile fatty acid solution (**Reagent #5**). Fill with $dH_2O$ to 750 mL (1.33X concentrated medium) and adjust pH to 7.5 (5 M NaOH). Attach a condenser to the Erlenmeyer flask and boil (10 min) to remove $O_2$ on a heating plate with a magnetic stir bar. Cool to ~60°C and add the remaining components (4 g $NaHCO_3$, 1 g L-cysteine HCl) under constant flushing with $CO_2$. After 10 min at ~60°C, fill the medium under constant $CO_2$ flushing in DURAN® Pressure plus flasks (size between 250-1000 mL) containing a magnetic stir bar. Close with airtight butyl stoppers and autoclave soon after. Store at room temperature.

*Note: 90 mL of 1.33X basal medium is required per 96-deepwell plate, when using 2 mL of medium per well and 60 wells per plate (outer wells filled with $dH_2O$;* ***see Step 4****).*

*To maintain fecal cultures similar to the original feces in terms of taxonomic composition, we suggest complementing the basal medium with the C-source mix, 6C+Muc (see heat-stable supplement: 15% starch, 15% pectin, 15% xylan, 8% arabinogalactan, 8% guar gum and 38% inulin and 10% mucin type II)***.**

**#8 Heat-stable supplement, *e.g.*, 6C+Muc (8X concentrated, anaerobic):** Dissolve 369 mg soluble starch, 369 mg pectin from citrus peel, 369 mg xylan, 185 mg arabinogalactan, 185 mg guar gum, 923 mg inulin and 240 mg mucin type II in 100 mL $dH_2O$ (resulting in an 8X concentrated solution) and adjust to pH 7. Attach a condenser to the Erlenmeyer flask and boil (10 min) to remove $O_2$. Cool under constant $N_2$ flushing. After 10 min at ~60°C, fill the solution under constant $N_2$ flushing in glass flasks, close with airtight butyl stoppers and autoclave soon after.

*Note: Heat-stable supplements can be flexibly adapted for testing other autoclavable components, e.g., reducing agents, xenobiotics, antibiotics or heat-stable vitamins. When calculating the final concentrations to use in the plate, consider that a concentrated stock solution should be prepared (in this protocol, supplements are 8X concentrated).*

**#9 Heat-sensitive supplement, *e.g.*, drug solutions (8X concentrated, anaerobic):** Dissolve the desired drug (8X concentrated) in anaerobic $dH_2O$ (**Reagent #11**) directly in the anaerobic chamber. Sterilize using a 0.2 µm syringe filter.

*Note: Heat-sensitive stock solution can be flexibly adapted. When calculating final concentrations, consider the higher concentration of the stock solution (here, 8X concentrated). When testing difficult-to-solubilize compounds, DMSO[1] can be used as a solvent instead (suggested final concentration of 0.2% in the plate).*

**#10 Phosphate buffer for diluting fecal samples (anaerobic):** Mix 190 mL mineral solution **I** (**Reagent #2**), 290 mL mineral solution **II** (**Reagent #3**) and 1 mL of resazurin stock solution (**Reagent #6**), adjust to pH 7 and fill up with $dH_2O$ to 1 L. Attach a condenser to the Erlenmeyer flask and boil (10 min) to remove $O_2$. Cool to ~60°C under constant $CO_2$ flushing and add the remaining components (4 g $NaHCO_3$, 1 g L-cysteine HCl). Flush with $CO_2$ until the solution turns transparent, fill the solution into Hungate tubes (9 mL per tube) under constant $CO_2$ flushing, close with airtight butyl stoppers and autoclave soon after.

**#11 Anaerobic $dH_2O$ (anaerobic):** Heat the desired volume of $dH_2O$ in a DURAN® Pressure plus flask under constant flushing with $N_2$. Boil (10 min), close with an airtight butyl stopper and autoclave soon after.

---

[1] *Chaveau, A., et al. (2023). Intestinal permeability and gut microbiota interactions of pharmacologically active compounds in valerian and St. John's wort. Biomed. Pharmacother.*

##### *C) Step-by-step procedure to cultivate gut microbiota/microbes in the anaerobic chamber using 96-deepwell plates*

**Step 1: Preparation of bYCFA and other solutions**

*Timing: At least two days before the experiment ~4 h*

Prepare reagents as described above, including bYCFA medium (**Reagent #7**; requires **Reagents #1-#5**), heat-stable supplement (*e.g.,* 6C+Muc, **Reagent #8**), heat-sensitive supplement (*e.g.,* drug solution, **Reagent #9**) and phosphate buffer (**Reagent #10;** requires **Reagents #2, #3 and #6**).

**Step 2: Preparation of the anaerobic chamber**

*Timing: One day before the experiment ~15 min*

The anaerobic chamber should have the following equipment: an incubator, a pH meter, multi-channel pipettes (*e.g.*, 100-1200 µL, 10-100 µL), a single-channel pipette (100-1000 µL), a pipette boy and a magnetic plate stirrer. To properly prepare the chamber, i) check gloves for holes, ii) replace the palladium catalysts with freshly-regenerated Stak-Paks (*i.e.*, incubation at 80 °C overnight), and iii) flush the chamber with gas mix containing $H_2$ (5% $H_2$, 10% $CO_2$, 85% $N_2$). Ideally, the experiment starts with $H_2$ above 2.5% (but below 5% due to explosion risks), as $H_2$ levels continuously decrease throughout the experiment. All material and reagents (needed for **Steps 3 to 8**), including autoclaved 96-deepwell plates (Ritter Medical), must be placed in the chamber at least 24 h before the experiment to allow proper removal of residual $O_2$. When bringing material into the chamber, puncture all packaging material with a needle to allow gas exchange.

**Step 3: Equilibration of bYCFA in the chamber**

*Timing: One day before the experiment ~10 min*

Place the bYCFA medium **(Reagent #7)** in the chamber at least 24 h before the experiment. Open the flask, cover with a Breathe-Easier Sealing Film (Diversified Biotech, Dedham, Massachusetts, United States), and place it on a stirrer to let the medium equilibrate with the gas composition of the chamber.

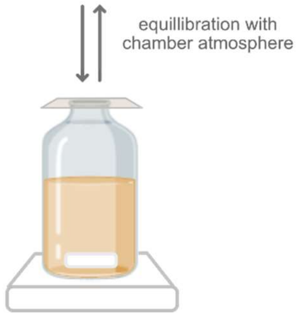


**Step 4: Preparation of the 96-deepwell plates with $dH_2O$ and compounds to test (*e.g.,* 6C+Muc and drug solutions)**

*Timing: Day of the experiment ~30-60 min*

Fill the outer wells of the 96-deepwell plate with sterile anaerobic $dH_2O$ to reduce evaporation of the media during the cultivation and prevent the edge effect[2]. Pipet 250 µL of 6C+Muc (heat-stable supplement; **Reagent #8**) and 250 µL of the drug solutions (heat-sensitive supplement; **Reagent #9**) or anaerobic $H_2O$ (control) into the desired wells using sterile containers and a multi-channel pipette.

*Note: The figure illustrates an exemplary plate layout to test two donors and nine drugs, which, however, can be flexibly adjusted.*

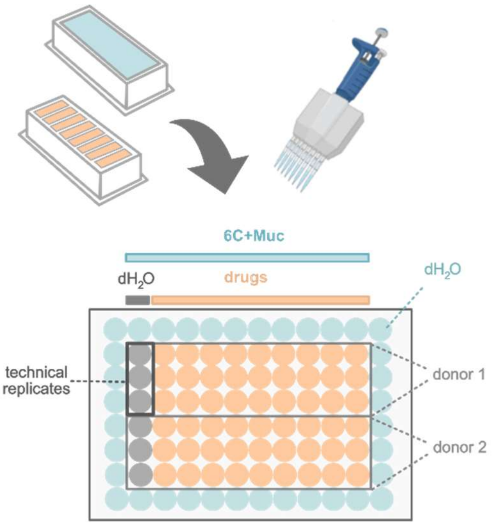


[2] *Douglas S. Auld, et al. (2020). Microplate Selection and Recommended Practices in High-throughput Screening and Quantitative Biology. Assay Guid. Man.*

#### Step 5: Preparation of the inoculum

Before inoculum preparation, sterilize the surfaces and gloves using a bleach solution (100 mL bleach, 60 mL of 5% acetic acid, 840 mL $dH_2O$) or sterilizing tissues (Wetrok Quick & Clean, Kloten, Switzerland).

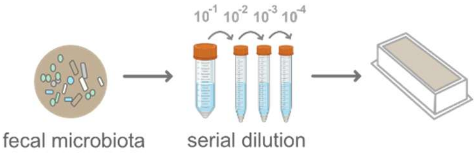


*Note: If the $O_2$ level rises while transferring the fecal sample or pure cultures into the chamber, wait until < 20 ppm.*

##### 5.1. Testing a fecal sample

*Timing: Day of the experiment ~15 min*

The fecal sample, collected in a plastic recipient containing an AnaeroGen™ 2.5L Sachet (Thermo Scientific™ Oxoid™, Waltham, MA, United States) to reduce the atmospheric oxygen, is brought into the chamber immediately upon arrival in the laboratory. For each donor microbiota, fill one 50 mL and three 15 mL Falcon tubes (for dilutions $10^{-1}$ to $10^{-4}$) containing 9 mL of phosphate buffer dilution solution (**Reagent #10**). Take ~1 g fecal sample using a sterile plastic spoon and resuspend it by gently shaking and mixing with the phosphate buffer solution ($10^{-1}$ dilution, 50mL Falcon tube). Let the main solid particles settle, then stepwise dilute down to the required dilution ($10^{-4}$) by transferring 1 mL into 9 mL phosphate buffer. Transfer the diluted fecal sample into a sterile container (*e.g.,* petri dish or reservoir). The fecal dilution is then ready to be inoculated.

##### 5.2. Testing pure bacterial cultures

*Timing: Day of the experiment ~15 min*

Typically, a pre-culture is prepared from a glycerol stock by 1-2% (v/v) inoculation of anaerobic bYCFA containing an appropriate C-source in Hungate tubes (37°C, 24 h). On the day of the experiment, $OD_{600}$ of the pre-culture is measured and if needed, the culture is diluted to a defined $OD_{600}$ value (*i.e.,* standardized inoculum). In the anaerobic chamber, transfer the culture into a sterile container (*e.g.,* petri dish or reservoir). Then, the culture is ready to be inoculated.

#### Step 6: Setting up starting pH of bYCFA and complementing media in 96-deepwell plates

*Timing: Day of the experiment ~5 min*

The pH meter should be calibrated on the day of the experiment (use new calibration solutions as the pH is altered when stored in the chamber). First, withdraw a small volume of equilibrated bYCFA medium (from **Step 3**) to measure pH (approximately 7.85). Next, calculate the amount of HCl needed to adjust the pH to the desired value using the formula **(2)**.

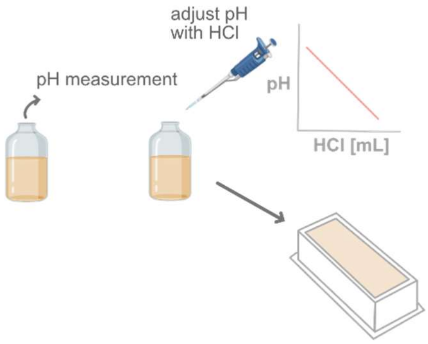


In brief, the calculations are based on a pre-determined calibration curve generated for bYCFA showing the change in pH upon HCl addition (as described in Figure 1E of the main article), resulting in the linear equation **(1)**.

$$pH = 7.85 - 45.1 * HCl_{add} \quad [mol/L] \qquad \textbf{(1)}$$

Using the slope of equation **(1)**, the amount of $HCL_{to\ add}$[mol/L] to add can be estimated (using equation **(2)**).

$$HCl_{to\ add}\ [mol/L] = \frac{pH_{final} - pH_{initial}}{40} \qquad \textbf{(2)}$$

After adding HCl, re-measure the pH and further adjust if needed. Then, fill 1.5 mL of bYCFA medium into each well of the 96-deepwell plate (pre-filled with supplements). Together with the previously pipetted supplements, the final total volume should be 2 mL.

**Step 7: Inoculation and incubation**

*Timing: Day of the experiment ~10 min*

Immediately inoculate the 96-deepwell plates with 20 µL of fresh inoculum (diluted fecal sample (**Step 5.1**) or pure culture (**Step 5.2**)) using a multi-channel pipette. Cover the plate with Breathe-Easier Sealing Film (Diversified Biotech) and incubate (37°C, 48 h).

**Step 8: Sampling and analysis.**

2 mL provides enough material for various types of analysis; i) $OD_{600}$ measurements by transferring the culture into a 96-well plate and measuring $OD_{600}$ using an absorbance plate reader (ca. 200 µL), ii) pH measurement by pipetting additional aliquots into well plates (ca. 200 µL) and measure pH (preferably performed in the anaerobic chamber) using a pH microelectrode (diameter 6 mm; Metrohm AG), iii) metabolite analysis by liquid chromatography using the supernatant (ca. 300 µL) and iv) 16S rRNA amplicon sequencing using cell pellet (ca. 1 to 2 mL).

To collect the supernatant and pellet, centrifuge cultures using a rotor adapted to 96-deepwell plates (5500 rpm, 20 min, 4°C). Then, split the pellet and resulting supernatants by pipetting the latter into another 96-deepwell plate. Both can be stored at -80°C (covered with an adhesive seal) until subsequent sample analysis. For organic acids quantification, *e.g.,* via liquid chromatography (HPLC or UPLC), supernatants are filtered using a 0.2 µm nylon membrane filter, preferably using a plate-based filtration system (AcroPrep™ Filter Plates; VWR International AG). The cell pellet can be resuspended in buffer for subsequent extraction of DNA.

Depending on the experimental design and the number of samples, pooling the technical replicates can be performed prior to analysis (see the Figure below). Though it is important to evaluate for each experimental set-up or sample type if pooling is reasonable and define criteria for pooling beforehand. Prepare two pools by transferring 0.5 mL from each replicate into a fresh plate (resulting in 1.5 mL sample volume). One pool can be used for direct analysis and the second sample pool can be stored at -80°C and considered as a backup.

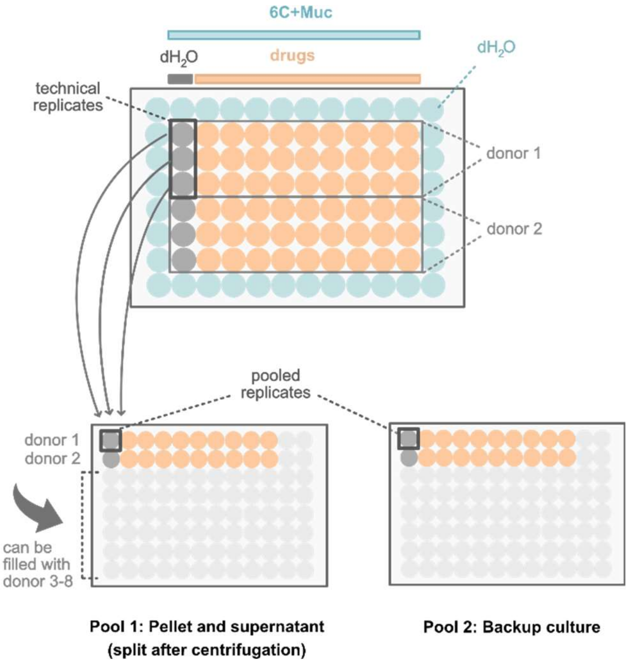
